# Seq-Scope: Submicrometer-resolution spatial transcriptomics for single cell and subcellular studies

**DOI:** 10.1101/2021.01.25.427807

**Authors:** Chun-Seok Cho, Jingyue Xi, Sung-Rye Park, Jer-En Hsu, Myungjin Kim, Goo Jun, Hyun-Min Kang, Jun Hee Lee

## Abstract

Spatial barcoding technologies have the potential to reveal histological details of transcriptomic profiles; however, they are currently limited by their low resolution. Here we report Seq-Scope, a spatial barcoding technology with a resolution almost comparable to an optical microscope. Seq-Scope is based on a solid-phase amplification of randomly barcoded single-molecule oligonucleotides using an Illumina sequencing-by-synthesis platform. The resulting clusters annotated with spatial coordinates are processed to expose RNA-capture moiety. These RNA-capturing barcoded clusters define the pixels of Seq-Scope that are approximately 0.5-1 μm apart from each other. From tissue sections, Seq-Scope visualizes spatial transcriptome heterogeneity at multiple histological scales, including tissue zonation according to the portal-central (liver), crypt-surface (colon) and inflammation-fibrosis (injured liver) axes, cellular components including single cell types and subtypes, and subcellular architectures of nucleus, cytoplasm and mitochondria. Seq-scope is quick, straightforward and easy-to-implement, and makes spatial single cell analysis accessible to a wide group of biomedical researchers.

## Introduction

The development of the light and electron microscopes profoundly contributed to the development of histology, the branch of biology which studies the microscopic anatomy of tissues (Mazzarini et al., 2020). Modern protein and mRNA detection techniques, such as immunohistochemistry and RNA in situ hybridization, further allowed for detecting the expression of specific biomolecules in histological slides (Callea et al., 1992). These technological advances tremendously strengthened our understanding of various physiological and pathophysiological processes and enabled the development of molecular diagnostic methods for various diseases.

Standard immunohistochemistry and RNA in situ hybridization can examine only one or a handful of target molecular species at a time; therefore, the amount of information obtained from a single experimental session is limited. To overcome this, emerging Spatial Transcriptomics (ST) techniques aim to examine all genes expressed from the genome from a single histological slide (Asp et al., 2020; Crosetto et al., 2015; Liao et al., 2020). There are three major methodologies of experimentally implementing ST. First, the sequential in situ hybridization method, often combined with combinatorial multiplexing, can increase the number of RNA species that can be detected from a single histological section. Second, in situ sequencing can identify RNA sequences from a tissue through fluorescence-based direct sequencing. Finally, the spatial barcoding method reveals both the RNA sequence and their spatial locations by capturing tissue RNA using a spatially-barcoded oligonucleotide array.

Among these three major methodologies, the spatial barcoding method is the most straightforward, comprehensive, widely-used and commercially available method easily accessible to many laboratories (Asp et al., 2020; Crosetto et al., 2015; Liao et al., 2020). Spatial barcoding is currently achieved by the microspotting of nucleotides (Salmen et al., 2018; Stahl et al., 2016), an array of split-pool-barcoded beads (Rodriques et al., 2019; Stickels et al., 2020; Vickovic et al., 2019), or a fabricated microfluidic channel (Liu et al., 2020). These methods, however, have an intrinsic limitation due to their low-resolution specifications. For instance, VISIUM from 10X Genomics, so far the only commercially available ST solution, has a center-to-center resolution of 100 μm (Bergenstrahle et al., 2020), which is worse than that of a naked eye (~40 μm). More recent technologies, such as Slide-Seq, HDST and DBiT-Seq, improved the resolution (Rodriques et al., 2019; Stickels et al., 2020; Vickovic et al., 2019); however, their resolution is still far inferior to optical microscope (0.5-1 μm). Consequently, detailed spatial organization of transcripts can only be inferred from pre-existing knowledge (Asp et al., 2019; Baccin et al., 2020; Bergenstråhle et al., 2020). Accordingly, none of the current spatial barcoding technologies have been able to reveal the microscopic details of the spatial transcriptome.

Here, we describe a technology for achieving sub-micrometer resolution spatial barcoding, designated as Sequence-Scope (Seq-Scope). Unlike former methods that use deterministic barcoding (Liu et al., 2020; Salmen et al., 2018; Stahl et al., 2016) or split-pool bead techniques (Rodriques et al., 2019; Stickels et al., 2020; Vickovic et al., 2019), our technique is based on the solid-phase amplification of a random barcode molecule, which can be conveniently achieved by the Illumina sequencing-by-synthesis (SBS) platform (Bentley et al., 2008). We show that this technology enables the generation of up to 1.5 million different spatially defined barcodes in a 1 mm^2^ area, which is substantially higher than any currently existing technologies by several orders of magnitude (e.g., ~10,000 fold over VISIUM (Bergenstrahle et al., 2020), ~600 fold over DBiT-seq (Liu et al., 2020), ~120 fold over Slide-Seq (Rodriques et al., 2019; Stickels et al., 2020) and ~15 fold over HDST (Vickovic et al., 2019)). Seq-Scope has a center-to-center resolution of 0.5-1 μm, far superior to previous technologies and comparable to an optical microscope. Using Seq-Scope, we obtained transcriptome images that clearly visualize microscopic cellular and subcellular structures of gastrointestinal tissues, such as the liver and colon, which were impossible to obtain through formerly existing methods.

## Results

### Seq-Scope Technology Overview

The Seq-Scope experiments are divided into two rounds of sequencing steps: 1^st^-Seq and 2^nd^-Seq (Fig. 1). 1^st^-Seq generates a physical array of spatially-barcoded RNA-capture molecules and a spatial map of barcodes where each barcoded sequence is associated with a spatial coordinate in the physical array. 2^nd^-Seq captures mRNAs released from the tissue placed on the physical array from the 1^st^-Seq, and sequences the captured molecules containing both cDNA and spatial barcode information.

**Fig 1.**
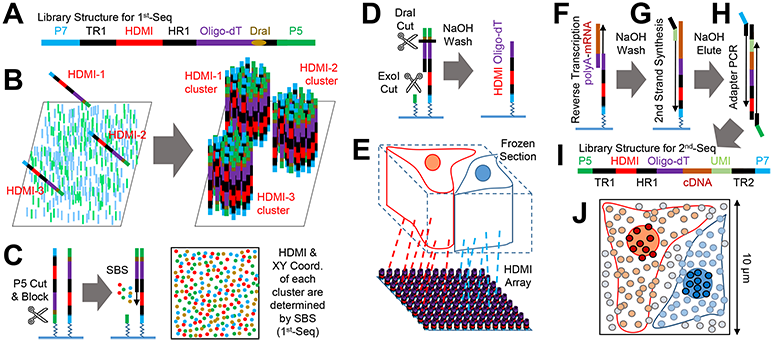
Seq-Scope Overview. (A) Schematic representation of the HDMI-oligo library structure. This library is used as an input of 1^st^-Seq, described below in (B) and (C). P5/P7, PCR adapters; TR1, TruSeq Read 1; HDMI, high-definition map coordinate identifier; HR1, HDMI Read 1. (B) Solid-phase amplification of different HDMI-oligo molecules on the flow cell surface. During 1^st^-Seq, a single “seed” molecule from the HDMI-oligo library forms a cluster of oligonucleotides that contain unique HDMI sequences. (C and D) Illumina sequencing by synthesis (SBS) determines the HDMI sequence and XY coordinates of each cluster (C). Then, HDMI oligonucleotide clusters are modified to expose oligo-dT, the RNA-capture domain (D). (E-I) HDMI-array captures RNA released from the overlying frozen section (E). Then, cDNA footprint is generated by reverse transcription of mRNA hybridized with oligo-dT domain (F). After that, secondary strand is synthesized using random priming method on the HDMI-cDNA chimeric molecule (G). Finally, adapter PCR (H) generates the sequencing library for 2^nd^-Seq (I), where paired-end sequencing using TR1 and TR2 reveals cDNA sequence and its matching HDMI barcode. TR2, TruSeq Read 2; UMI, unique molecule identifier. (J) HDMI-array contains up to 150 HDMI clusters in 100 μm^2^ area. Each cluster has over 1,000 RNA capture probes with unique HDMI sequences.

1^st^-Seq of Seq-Scope starts with the solid-phase amplification of a single-stranded synthetic oligonucleotide library using an Illumina SBS platform (MiSeq in the current study; Fig. 1A). The oligonucleotide “seed" molecule contains the PCR/read adapter sequences, the restriction enzyme-cleavable RNA-capture domain (oligo-dT), and most importantly, the high-definition map coordinate identifier (HDMI), a spatial barcode composed of a 20-32 random nucleotide sequence. The “seed” oligonucleotide library is amplified on a lawn surface coated with PCR adapters (Fig. 1B), generating a number of clusters, each of which are derived from a single “seed” molecule. Each cluster has thousands of oligonucleotides that are identical clones of the initial oligonucleotide “seed” (Bentley et al., 2008) (Fig. 1B). The HDMI sequence and spatial coordinate of each cluster are determined through a typical SBS procedure using the MiSeq Control Software (MCS) equipped with Real-Time Analysis (RTA) (Fig. 1C and S1A) (Ravi et al., 2018). After SBS, oligonucleotides in each cluster are processed to expose the nucleotide-capture domain (Fig. 1D and S1A), producing an HDMI-encoded RNA-capturing array (HDMI-array; Fig. 1E), the physical array produced by 1^st^-Seq of Seq-Scope.

2^nd^-Seq of Seq-Scope begins with overlaying the tissue section slice onto the HDMI-array (Fig. 1E). The mRNAs from the tissue are used as a template to generate cDNA footprints on the HDMI-barcoded RNA capture molecule (Fig. 1F and S1B). Then the secondary strand is synthesized on the cDNA footprint using an adapter-tagged random primer (Fig. 1G and S1B). Since each cDNA footprint is paired with a single random primer after washing, the random priming sequence is used as a unique molecular identifier (UMI; Fig. S1B). The secondary strand, which is a chimeric molecule of HDMI and cDNA sequences, is then collected and prepared as a library through PCR (Fig. 1H and S1B). The paired-end sequencing of this library reveals the cDNA footprint sequence, as well as its corresponding HDMI sequence (Fig. 1I and S1B).

For each HDMI sequence, 1^st^-Seq provides spatial coordinate information while 2^nd^-Seq provides captured cDNA information. Correspondingly, the spatial digital gene expression (DGE) matrix is constructed by combining the 1^st^-Seq and 2^nd^-Seq data (Fig. S1C-S1E). The spatial DGE matrix is used for various analyses, including gene expression visualization and clustering assays (Fig. S1C-S1E).

### HDMI-Array Captures Spatial RNA Footprint of Tissues

Through a series of titration experiments (Fig. S2A and S2B), we produced the HDMI-array with a sequenced cluster density of up to 1.5 million clusters per mm^2^ (Fig. 2A, S2A and S2B). The distance between the centers of nearby clusters was estimated to be between 0.5-1 μm (Fig. 2A and S2A). Seq-Scope generates up to 150 HDMIs in a 100 μm^2^ area; this resolution should be sufficient to perform single cell and subcellular analysis of spatial transcriptome (Fig. 1J).

**Fig 2.**
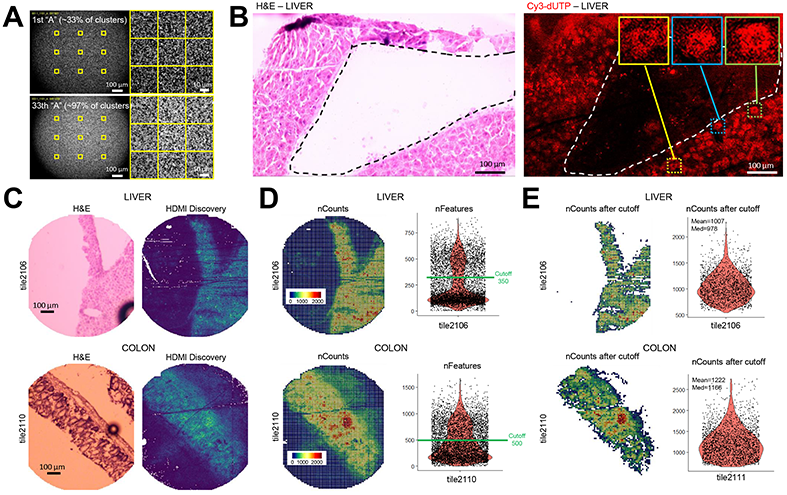
Seq-Scope Captures and Analyzes Spatial cDNA Footprint from Tissue-Derived RNA. (A) Representative images of HDMI clusters in the HDMI-array, retrieved from the Illumina sequence analysis viewer. Upper panel visualizes “A” intensity at the 1^st^ cycle of the 1^st^-Seq SBS, where 33% of HDMI clusters exhibit fluorescence. Lower panel visualizes “A” at the 33^rd^ cycle, where 97% of HDMI clusters exhibit fluorescence. Yellow squares in the left panels are magnified in the right panels. (B) H&E staining and its corresponding Cy3-dUTP labeling fluorescence images from fragmented liver section. Gross tissue boundaries (dotted lines) are well preserved in the underlying cDNA footprint. Box insets in the right panel highlights single cell-like patterns in the cDNA footprint. (C) H&E staining and its corresponding HDMI discovery plot drawn from the analysis of 1^st^-Seq and 2^nd^-Seq outputs. Brighter color in the HDMI discovery plot indicates that more HDMI was found from 2^nd^-Seq in the corresponding pixel area. (D and E) After data binning with 10 μm-sided square grids, number of UMI counts (D, left; nCounts) and gene features (D, right; nFeatures) of liver (upper) and colon (lower) datasets were presented in spatial and violin plot, respectively. Setting a 350 (liver) or 500 (colon) cutoff was sufficient to isolate grids covered by the tissue area (E, left), each of which contains approximately 1,000 UMIs (E, right).

The RNA-capturing capability of the HDMI-array was first evaluated by performing a Cy3-dUTP-mediated cDNA labeling assay using a fragmented frozen liver section. The HDMI-array successfully captured tissue transcriptome and generated a spatial cDNA footprint that preserved the gross shape of the overlying tissue (Fig. 2B). Interestingly, the Cy3-dUTP labeling assay also revealed microscopic details of cDNA footprints that resemble a single cell morphology (Fig. 2B, insets), which has a fluorescence texture that is similar to those produced by the underlying clusters (Fig. 2A and S2A).

Then we performed the full Seq-Scope procedure (1^st^-Seq and 2^nd^-Seq; Fig. 1) on two representative gastrointestinal tissues, liver and colon. In each 1^st^-Seq experiment, the HDMI-array was produced in 1 mm-wide circular areas of the MiSeq flow cell, also known as “tiles” (Bentley et al., 2008; Ravi et al., 2018) (tile ID: 2101 to 2119; Fig. S2C). Liver and colon tissue sections were overlaid onto the HDMI arrays, examined by H&E staining, and subjected to 2^nd^-Seq. Analysis of the 1^st^-Seq and 2^nd^-Seq data (Fig. S1C) demonstrated that the RNA footprints were discovered mostly from tissue-overlaid regions (Fig. 2C, S2D and S2E), confirming that Seq-Scope can indeed capture and analyze the spatial transcriptome from the tissues.

### Seq-Scope Captures Transcriptome Information with High Efficiency

Although each HDMI-barcoded cluster covers an extremely tiny area (less than 1 μm^2^), many HDMI clusters were able to identify 10-100 unique transcripts from the overlying tissue section (Fig. S2F and S2G). To compare the data output of Seq-Scope with other existing ST technologies, we quantified the number of gene features and unique transcripts in 10 μm-sided square grids (Fig. S2H and S2I). Since the tissue-overlaid grids distinctively displayed a higher number of gene features and unique transcript counts (Fig. 2D, S2H and S2I), setting a simple gene feature cutoff was sufficient to isolate tissue-overlaid grids (Fig. 2E, S2J-S2M). The average number of unique transcripts per grid varied substantially from tile to tile (Fig. S2N and S2O), but the tiles located at the center of the MiSeq flow cell revealed up to 1,000-1,200 average unique transcripts per 100 μm^2^ grid area (Fig. 2E). Considering that the current data are estimated to represent only ~70% (liver) and ~42% (colon) of the total library complexity (Fig. S2P), the maximum possible Seq-Scope output could be estimated to be over 2,000 UMI per 100 μm^2^. The number of genes discovered from each Seq-Scope session is 25,180 (liver) and 24,927 (colon). All these output measures are superior or comparable to most existing spatial barcoding technologies, as discussed in more detail in the Discussion. Therefore, Seq-Scope provides an outstanding mRNA capture output, in addition to providing an unmatched spatial resolution output.

### Seq-Scope Reveals Nuclear-Cytoplasmic Transcriptome Architecture from Tissue Sections

mRNA is transcribed and poly-A modified in the nucleus. Before it can be transported to the cytoplasm, it is spliced, and intronic sequences are removed. Therefore, the nuclear area has a higher concentration of unspliced mRNA sequences, while the cytoplasmic area has a higher concentration of spliced mRNA sequences (Fig. 3A). Several RNAs in mouse liver, such as Malat1, Neat1 and Mlxipl, exhibit nuclear localization with strong attenuation in cytoplasmic transport (Fig. 3A) (Bahar Halpern et al., 2015). On the other hand, mitochondria in the cytoplasm has a unique transcriptome structure with mitochondria-encoded RNAs (mt-RNA; Fig. 3A).

**Fig 3.**
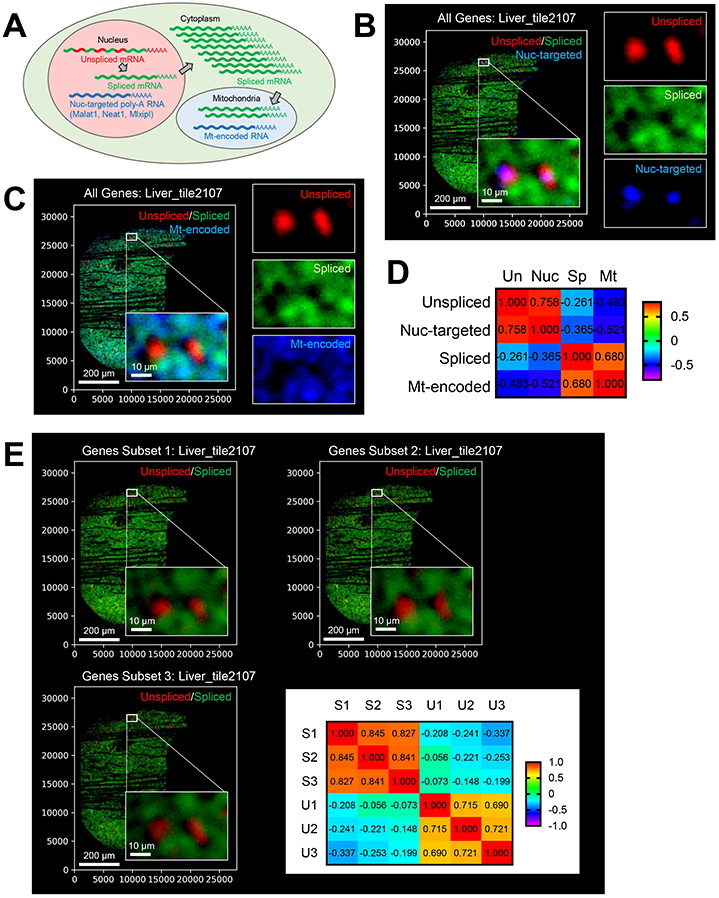
Seq-Scope Visualizes Subcellular Spatial Transcriptome. (A) Schematic diagram depicting the distribution of different RNA species in subcellular compartments. (B-D) Spatial plot of all unspliced and spliced transcripts, as well as RNA species that are known to localize to nucleus (Nuc-targeted; *Malat1, Neat1* and *Mlxipl*) in liver (B). RNA species that are encoded by mitochondrial genome (Mt-encoded) were also analyzed (C). Pearson correlations (r) between these transcript intensities were presented as a heat map (D). (E) Spatial plot of unspliced and spliced transcripts in three independent subsets of genes (Gene Subset 1-3). Pearson correlations (*r*) between these transcript intensities were presented as a heat map. S1-3, Spliced 1-3; U1-3, Unspliced 1-3.

To examine the subcellular-level spatial transcriptome through Seq-Scope, we plotted all spliced and unspliced transcripts discovered from Seq-Scope in a two-dimensional histological coordinate plane. Intriguingly, unspliced transcripts showed an interesting pattern as their expression was restricted in tiny circles with a diameter of approximately 10 μm, which is about the size of hepatocellular nuclei (Baratta et al., 2009) (Fig. 3B and S3A). More interestingly, spliced mRNAs were less frequently discovered in the unspliced area, while nuclear-localized RNAs, including Malat1, Neat1 and Mlxipl (Bahar Halpern et al., 2015), were more frequently found in the unspliced area (Fig. 3B). On the other hand, mt-RNAs were more frequently found in the spliced cytoplasmic area (Fig. 3C and S3B). As a result, quantification of single cell images revealed strong positive correlations between unspliced and nuclear-localized mRNAs and between spliced and mitochondrial mRNAs, while displaying strongly negative correlations between the opposite groups (Fig. 3D and S3C).

These results suggest that the relative proportion of spliced and unspliced transcripts can be useful metrics to determine the nuclear-cytoplasmic structure from the Seq-scope dataset. To further test the robustness of these observations, we randomly divided all genes into three independent subsets, calculated expressions of spliced and unspliced mRNAs from each gene subset, and analyzed each dataset through the same plotting method. All three datasets similarly visualized a nuclear-cytoplasmic structure with a strong statistical correlation (Fig. 3E and S3D). Therefore, as predicted (Fig. 1J), Seq-Scope can visualize the nuclear-cytoplasmic transcriptome structure from the tissue sections.

### Seq-Scope Reveals Spatial Transcriptomic Details of Metabolic Liver Zonation

To systematically characterize the heterogeneity of the liver cell transcriptome, we analyzed the square-gridded Seq-Scope dataset (Fig. S2K and S2N) with the standard scRNA-seq analysis pipeline (Stuart et al., 2019). Multi-dimensional clustering analysis identified many interesting and biologically-relevant cell types (Fig. S4A) with a long list of cluster-specific marker genes (Fig. S4B-S4D and Table S1).

Hepatocytes, the parenchymal cell type of liver, are exposed to varying gradients of oxygen and nutrients according to their histological locations, leading to metabolic zonation whereby cells express different genes to perform their zone-specific metabolic functions (Zone 1-3 or Z1-3; Fig. 4A) (Ben-Moshe and Itzkovitz, 2019). Zone 1 marks periportal hepatocytes close to the hepatic portal vein and artery, while zone 3 marks pericentral hepatocytes close to the central vein (Fig. 4A). Zone 2 are intermediate hepatocytes located between zones 1 and 3 (Fig. 4A).

**Fig 4.**
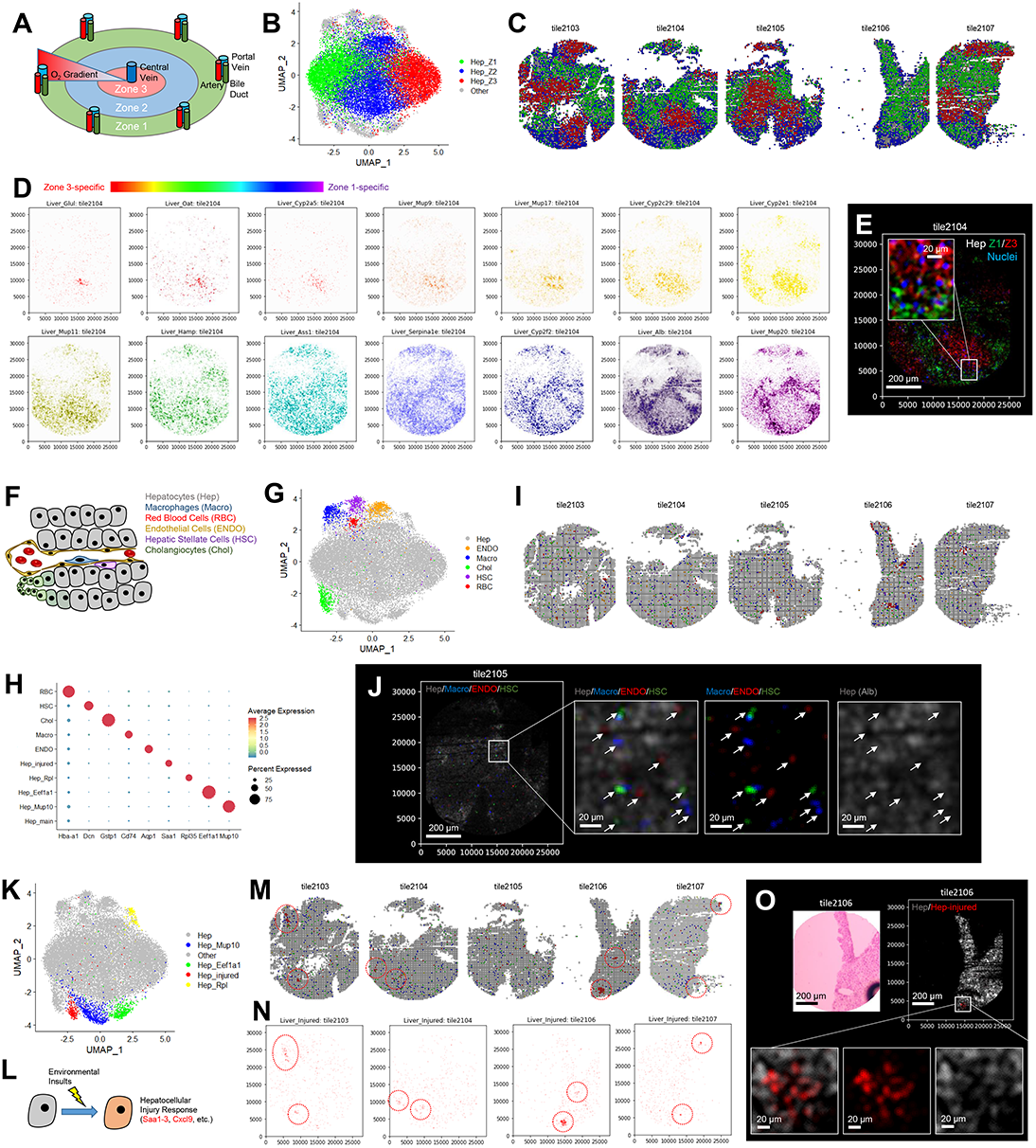
Seq-Scope Reveals Microscopic Details of Spatial Transcriptome in Normal Liver. Spatial transcriptome of normal liver was analyzed using Seq-Scope (tile ID: 2102-2107). (A-C) Schematic diagram depicting metabolic zonation of normal liver (A). UMAP (B) and spatial plots (C) visualize clusters of 10 μm-sided square grid units representing zonated hepatocytes in the indicated color. (D) Spectrum of genes exhibiting different zone-specific expression patterns were examined by spatial plot analysis. Zone 3-specific genes are depicted in warm (red-orange-yellow) colors, while zone 1-specific genes are depicted in cold (blue-purple) colors. (E) Zone 1-specific transcripts (green), zone 3-specific transcripts (red) and unspliced transcripts (blue, Nuclei) were visualized together in a histological coordinate plane. (F-J) Schematic diagram depicting cellular components of normal liver (F). UMAP (G) and spatial plots (I) visualizing clusters of 10 μm-sided square grid units representing indicated cell types. Cell type-specific transcript expression was analyzed using dot plot (H) and gene expression plot (J) analyses. (K-O) UMAP (K) and spatial plots (M) visualizing clusters of grid pixels representing indicated hepatocellular subpopulations. Marker genes for Hep_injury cluster, which represent transcriptome responses to hepatocellular injury (L), were examined through spatial gene expression plotting (N and O).

Multi-dimensional clustering analysis identified zonated hepatocytes as the major clusters found from the Seq-Scope dataset (Fig. 4B and S4A). Spatial plotting of the cluster identity clearly visualized zones 1-3 in the two-dimensional grid space (Fig. 4C). To fully utilize the submicrometer resolution performance of Seq-Scope, we directly plotted zone-specific molecular markers into the histological coordinate plane. Direct plotting analysis revealed a spectrum of genes showing various zonation patterns, which cannot be explained by the three simple layers (Fig. 4D). For instance, the immediate pericentral hepatocytes specifically expressed extreme zone 3 markers such as *Glul* and *Oat* (Fig. 4D, reds). *Cyp2a5, Mup9* and *Mup17* were also narrowly expressed in hepatocytes surrounding the central vein; however, *Mup9* and *Mup17* displayed lower expression in the immediate pericentral hepatocytes, forming a small donut-like staining pattern (Fig. 4D, oranges). In contrast, general pericentral markers, such as *Cyp2c29* and *Cyp2e1*, were broadly expressed across all pericentral hepatocytes (Fig. 4D, yellows). Several genes, such as *Mup11* and *Hamp*, were not expressed in extreme zone 1 and zone 3 layers but showed higher expression in the intermediary layers (Fig. 4D, greens). Likewise, different periportal markers, such as *Ass1, Serpina1e, Cyp2f2, Alb and Mup20*, exhibited various levels of zone 1-specific expression patterns (Fig. 4D, blues and purples). Many of these observations are supported by previous scRNA-seq, RNA *in situ* hybridization (Aizarani et al., 2019; Halpern et al., 2017) and immunostaining results (Park et al., 2020).

Interestingly, most of these zone 1- or zone 3-specific markers were cytosolically located, as they did not overlap with the unspliced transcript-enriched area (Fig. 4E). This finding is consistent with the notion that zone-specific proteins are actively translated in the cytosol to perform zonated metabolic functions (Aizarani et al., 2019; Ben-Moshe and Itzkovitz, 2019; Halpern et al., 2017; Park et al., 2020). In contrast to zone 1 and zone 3 hepatocytes that exhibit clear transcriptome features, zone 2 hepatocytes, which do not exhibit obvious periportal or pericentral transcriptome characteristics, were subclustered based on their subcellular transcriptome heterogeneity. Correspondingly, Zone 2 hepatocytes were found in clusters enriched with nuclear transcripts (*Malat1, Neat1* and *Mlxipl*; cluster 9 in Fig. S4A), mitochondrial transcripts (mtRNA; cluster 1 in Fig. S4A) and long non-coding RNAs (lncRNA; cluster 3 in Fig. S4A) with heterogeneous spatial gene expression patterns (Fig. S4E).

### Seq-scope Detects Non-Parenchymal Cell Transcriptome from Liver Section

Although hepatocytes are the major cellular component in the liver, several non-parenchymal cells, such as macrophages (Macro), hepatic stellate cells (HSC), endothelial cells (ENDO), cholangiocytes (Chol), as well as red blood cells (RBC), can be found in a small portion of the histological area (Fig. 4F) (Ben-Moshe and Itzkovitz, 2019). Multi-dimensional clustering analysis of the gridded Seq-Scope data identified clusters that correspond to these non-parenchymal cell types (Fig. 4G), based on expression of macrophage markers (*Clec4f, Cd74, Cd5l*, MHC-II and complement components), HSC markers (*Dcn* and *Ecm1*; (Xiong et al., 2019)), sinusoidal endothelial markers (*Aqp1, Dnase1l3, Kdr, Oit3* and *Stab2*; (de Haan et al., 2020)), cholangiocyte/oval cell markers (*Gstp1* and *Gstp2*; (Tee et al., 1996)) and RBC markers (hemoglobins) (Fig 4H, S4B and S4D). Although most of the histological space was occupied by the hepatocellular area, the small, fragmented spaces scattered throughout the section represented the non-parenchymal cell area (Fig. 4I). Spatial plotting onto the histological coordinate plane indeed demonstrated that hepatocyte marker Alb expression is sharply diminished where macrophage, HSC and endothelial cell markers are abundantly expressed (Fig. 4J), consistent with the knowledge that hepatocyte markers are scarcely expressed in non-parenchymal cells (Werner et al., 2015).

### Seq-scope Identifies Hepatocyte Subpopulations undergoing Tissue Injury Response

Clustering also identified minor hepatocyte subpopulations (Fig. 4K) expressing hepatocyte injury response genes (*Saa1-3* and *Cxcl9*; Fig. 4L) (Sack, 2020; Saiman and Friedman, 2012), a subset of major urinary proteins (*Mup10, Mup14* and *Mup7*), a translation elongation factor (*Eef1a1*) that was formerly associated with hepatocarcinogenesis (Abbas et al., 2015), and a subset of ribosomal proteins (*Rpl15, Rpl35* and their matching pseudogenes) (Fig. 4H). These clusters were spatially scattered throughout the liver sections (Fig. 4M and S4F), although the cluster expressing injury response markers (Fig. 4L) was restricted to a localized area (dotted red circles in Fig. 4M and 4N). In the spatial plotting analysis, expression of the liver injury markers substantially overlapped with *Alb* (Fig. 4O), confirming that they are hepatocyte subpopulations with an altered transcriptome.

Processing the normal liver Seq-Scope data through smaller grids, including 7 μm (Fig. S4G-S4L) and 5 μm (Fig. S4M-S4R) square grids, also robustly identified hepatocyte zonation, parenchymal/non-parenchymal cells and hepatocyte subpopulations, confirming that our observations described here are significant and reproducible.

### Seq-Scope Reveals Transcriptomic Details of Histopathology Associated with Liver Injury

The data presented above confirm that Seq-Scope can reveal the transcriptome heterogeneity and spatial complexity of the normal liver at various scales. But could Seq-Scope also reveal the pathological details of transcriptome dysregulation in diseased liver? To address this, we utilized our recently developed mouse model of early-onset liver failure that was provoked by excessive mTORC1 signaling (Cho et al., 2019). This model (*Tsc1*^*Δhep*^*/Depdc5*^*Δhep*^ mice or *TD* mice) is characterized by a widespread hepatocellular oxidative stress, leading to localized liver damage, inflammation and fibrotic responses (Cho et al., 2019).

We first examined the cellular components of the *TD* liver through clustering analysis of the grid pixel dataset. Most cell types identified from the normal liver, such as zone 1/2/3 hepatocytes, macrophages, HSCs, endothelial cells, and RBC, were also discovered from the *TD* liver (Fig. S5A-S5C; Table S1). As observed from the normal liver dataset, some zone 2 hepatocyte clusters represented nuclear and mitochondrial areas (Fig. S5A), and nuclear, cytoplasmic and mitochondrial structures were clearly visualized through unspliced, spliced and mitochondria-encoded transcripts, respectively (Fig. S5D).

Former bulk RNA-seq results showed that the *TD* liver upregulates oxidative stress signaling pathways. Consistent with this, Seq-Scope identified that the *TD* liver expresses elevated levels of several antioxidant genes such as *Gpx3* and *Sepp1.* Interestingly, induction of these genes was very strong in zone 1 hepatocytes, while the upregulation was not pronounced in zone 3 hepatocytes (Fig. S5E). Therefore, the oxidative stress response of the *TD* liver is zone-specific.

We focused on the cell types involved in the liver injury response, such as macrophages and HSCs, as well as hepatocytes expressing injury response markers (Fig. 5A). Notably, hepatocytes exhibiting tissue injury responses (Hep_injured) were much more prevalent in the *TD* liver, and a novel type of injured hepatocyte subpopulation expressing genes encoding clusterin, MMP-7 and osteopontin proteins (Hep_novel) was observed (Fig.5A and S5F; Table S1). These altered hepatocytes, as well as macrophages and HSCs, were located in an area where gross liver injury and inflammatory features were observed from the H&E histology analysis (Fig. 5B and 5C, dotted rectangles). At the site of the liver injury, the dead cell area (yellow stars in Fig. 5C and 5D, where RNA footprint or cytoplasmic eosin staining was not discovered) was surrounded by a layer of inflamed macrophages and then by a layer of hepatocyte subpopulations that consisted of Hep_injured and Hep_novel (Fig. 5B-5D). HSCs also infiltrated into the macrophage layers (Fig. 5E). As expected, expression of *Alb*, the hepatocyte marker, was found in the area occupied by Hep_injured and Hep_novel populations but not by macrophage or HSC populations (Fig. 5D and 5E).

**Fig 5.**
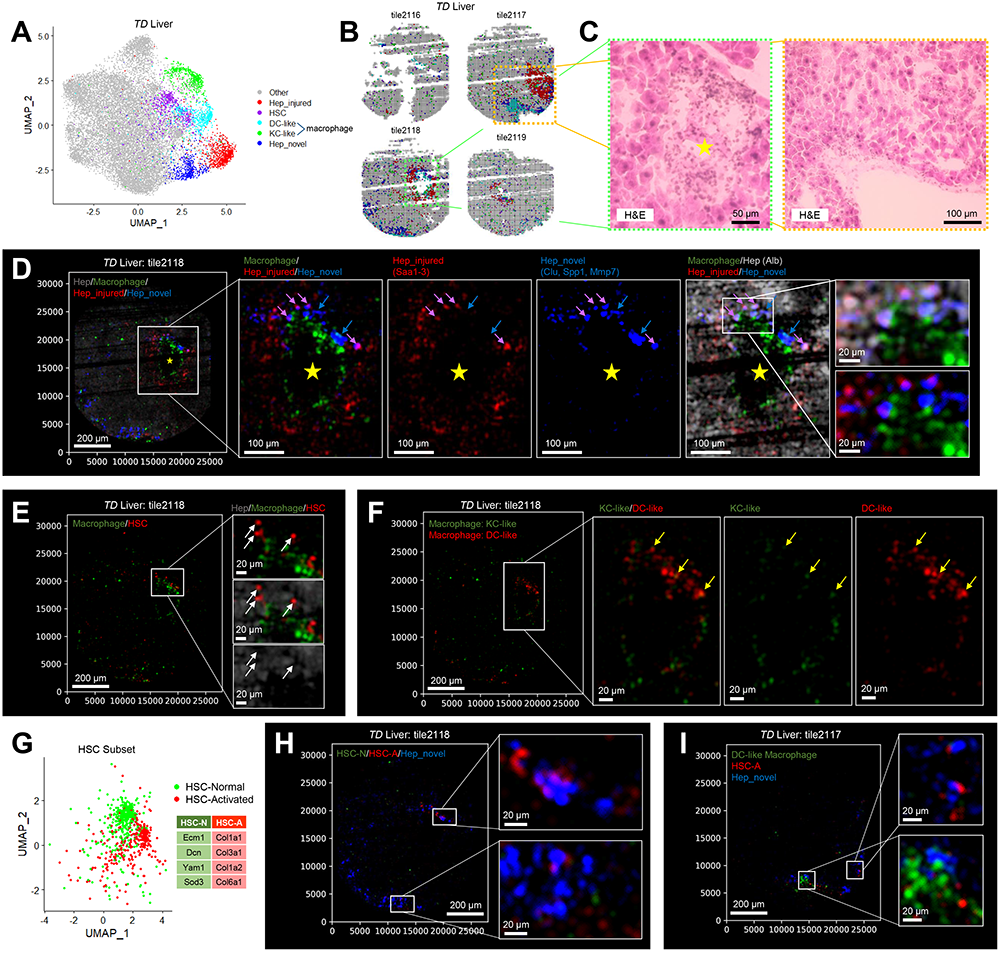
Seq-Scope Examines Liver Histopathology at Microscopic and Transcriptomic Scales. Liver from a *Tsc1*^*Δhep*^/*Depdc5*^*Δhep*^ (*TD*) mouse, which suffers liver injury and inflammation (Cho et al., 2019), was examined through Seq-Scope (tile ID: 2116-2119). (A-C and G) UMAP (A) and spatial plots (B) visualize clusters of 10 μm-sided square grid units representing injury-responsive cell populations. (C) H&E images correspond to the boxed regions in (B). Yellow star in (C) marks the area of hepatocellular injury. (G) Normal (HSC-N) and activated (HSC-A) subclusters of HSC population were identified, and their marker genes are listed in a table. (D-F, H and I) Marker genes for indicated cell types were plotted onto the histological coordinate plane with indicated colors. Boxed areas are magnified on right. Yellow stars in (D) indicate dead cell area where RNA footprint was scarcely discovered. Blue arrows in (D) indicate hepatocytes expressing Hep_novel markers. Purple arrows in (D) indicate hepatocytes expressing both Hep_injured and Hep_novel markers. In (D), Hep_novel and Hep_injured markers overlap with *Alb*, the hepatocyte marker, while macrophage markers do not overlap with *Alb* and other hepatocyte markers. White arrows in (E) indicate the position of cells expressing HSC markers but not hepatocyte marker *Alb*. Yellow arrows in (F) indicate macrophages expressing both KC-like and DC-like markers.

In the *TD* liver dataset, the number of macrophage populations was increased, and the two subgroups of macrophages, dendritic cell-like (DC-like) and Kupffer cell-like (KC-like) populations, were identified. DC-like cells expressed dendritic cell markers such as *Cd74* and MHC-II component genes, while KC-like cells expressed Kupffer cell-specific markers such as *Clec4f* (Fig. S5F; Table S1). Most macrophages expressed either DC-like or KC-like markers at the liver injury site, while a smaller number of cells expressed both markers (yellow arrows in Fig. 5F). Similarly, the HSC population was also increased and differentiated into normal HSC (HSC-N) and activated HSC (HSC-A) subpopulations (Fig. 5G); HSC-A exhibited fibroblast-like characteristics such as high collagen expression (Fig. 5G). Interestingly, DC-like macrophages and HSC-A were frequently found in the area where Hep_novel cells were discovered (Fig. 5D, 5F, 5H and 5I), suggesting that these cell types are specifically involved in fibrotic responses to liver injury.

Although gene expression markers for DC-like macrophages, HSC-A, Hep_injured and Hep_novel populations showed prominent localization pattern around the injury site (Fig. S5G-S5J), markers for KC-like macrophages and HSC-N populations were more equitably distributed throughout the *TD* liver section (Fig. S5K and S5L), suggesting that KC-like and HSC-N populations play a more homeostatic role over the inflammatory role.

### Seq-Scope Visualizes Histological Layers of Colon Tissue

The colon is another gastrointestinal organ with complex tissue layers, histological zonation structure, and diverse cellular components (Levine and Haggitt, 1989). Using the colon, we examined whether Seq-Scope can examine spatial transcriptome in a non-hepatic tissue.

The colonic wall can be histologically divided into the colonic mucosa and the external muscle layers (Farkas et al., 2015). The colonic mucosa consists of the epithelial layer, lamina propria and muscularis mucosae (M. mucosae), a very thin layer of smooth muscle that separates mucosa from underlying submucosa and smooth muscle layers (Fig. 6A). The epithelial layer is further divided into crypt-base, transitional and surface layers (Fig. 6A) (Farkas et al., 2015). Multi-dimensional clustering analysis of the gridded Seq-Scope dataset of the colon slice (Fig. S6A, S6B and Table S2) revealed these layers as the major clusters of the colonic transcriptome (Fig. 6B): epithelial layers of crypt base, transitional and surface cells, as well as lamina propria and smooth muscle layers, were identified from the grids. The spatial plotting assays visualized histological layers on the gridded plane (Fig. 6C and S6C) or histological coordinate plane (Fig. 7A and S7A). Although M. mucosae was not identifiable from the gridded dataset (Fig. 6C and S6C), direct plotting onto the histological coordinate plane enabled its visualization (white arrows in Fig. 7A), indicating that Seq-Scope can detect all the major histological layers of the colon tissue.

**Fig 6.**
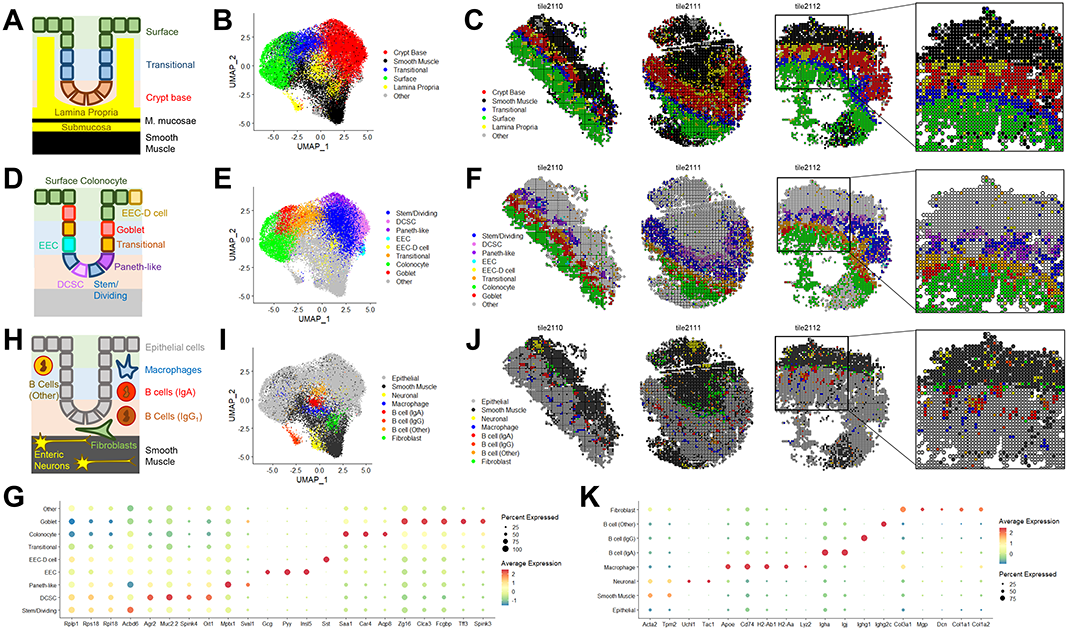
Seq-Scope Identifies Various Cell Types from Colonic Wall Histology. Spatial transcriptome of colon was analyzed using Seq-Scope. Seq-Scope dataset processed with 10 μm-sided square gridding was analyzed. (A-C) Seq-Scope reveals major histological layers through transcriptome clustering. (A) Schematic representation of colonic histology. Clusters corresponding to the indicated histological layers were visualized in UMAP manifold (B) and histological space (C). (D-G) Seq-Scope reveals epithelial cell diversity through transcriptome clustering. (D) Schematic representation of colonic epithelial cell types. Clusters corresponding to the indicated epithelial cell types were visualized in UMAP manifold (E) and histological space (F). Cluster-specific markers were examined in dot plot analysis (G). (H-K) Seq-Scope reveals non-epithelial cell diversity through transcriptome clustering. (H) Schematic representation of colonic non-epithelial cell types. Clusters corresponding to the indicated non-epithelial cell types were visualized in UMAP manifold (I) and histological space (J). Cluster-specific markers were examined in dot plot analysis (K).

**Fig 7.**
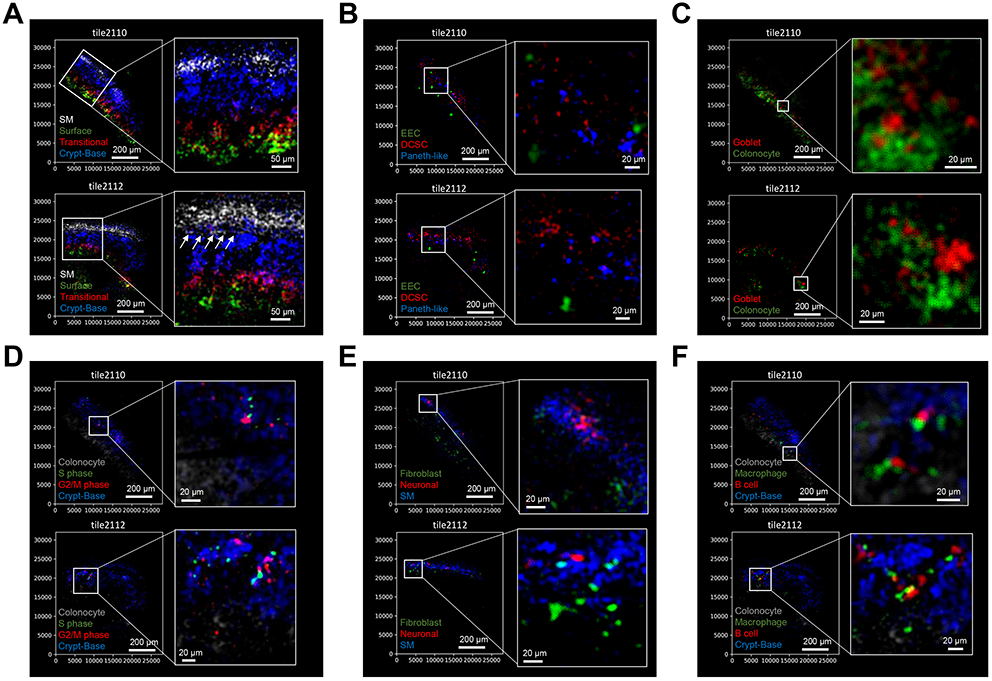
Seq-Scope Enables Microscopic Analysis of Colon Spatial Transcriptome. Colonic spatial transcriptome was analyzed using Seq-Scope. Original Seq-Scope dataset was analyzed by gene expression plotting on the histological coordinate plane, using cell type-specific marker genes. (A) Histological layers of smooth muscle, crypt base, transitional and surface epithelial cells were visualized through spatial gene expression plot. White arrows in (A) indicate the position of muscularis mucosae, the thin layer of smooth muscle separating mucosa and submucosa. (B and C) Different types of epithelial cells, including enteroendocrine cells (EEC), deep crypt secretory cells (DCSC), Paneth-like cells, Goblet cells and colonocytes were visualized through spatial plotting of cell type-specific marker genes. (D) Expression of S phase- and G2/M phase-specific marker genes were plotted onto the histological coordinate plane. Colonocyte and crypt base marker genes were also plotted to provide a background reference. (E and F) Different types of non-epithelial cells were visualized through spatial plotting of cluster-specific marker genes. In (F), colonocyte and crypt base marker genes were also plotted to provide a background reference.

### Seq-Scope Identifies Individual Cellular Components from Colonic Epithelia

We took a closer look into the colonic epithelial cell types (Fig. 6D). Cells in the crypt base were divided into three subgroups: Stem/dividing cells, deep crypt secretory cells (DCSC) and Paneth-like cells (Fig. 6E and 6F). The stem/dividing cells expressed higher levels of ribosomal proteins while lacking markers expressed in the other epithelial cell types (Fig. 6G; Table S2); these results are consistent with former scRNA-seq (Haber et al., 2017) and laser-capture microdissection (Moor et al., 2018) studies of intestinal single cell diversity. In contrast, DCSCs (Altmann, 1983; Sasaki et al., 2016), also known as colonic crypt base secretory cells (Rothenberg et al., 2012), pregoblet cells (Park et al., 2009) or crypt-located goblet cells (Parikh et al., 2019), expressed high levels of secretory cell markers, such as *Agr2, Muc2, Spink4* and *Oit1* (Fig. 6G; Table S2). Interestingly, a cell type with high expression of the mucosal pentraxin precursor (*Mptx1*), a recently identified marker of the Paneth cell in the small intestine (Haber et al., 2017), was discovered from the Seq-Scope results. Although the Paneth cell is largely absent from the colon and Paneth cell function is partially mediated by DCSCs (Rothenberg et al., 2012; Sasaki et al., 2016), this result suggests that the colon may possess a cell type whose transcriptome and function is more similar to the Paneth cell (Paneth-like cell). The expression of DCSC and Paneth-like cell markers showed non-overlapping patterns of expression in the histological coordinate plane (Fig. 7B, S7B and S7C) and the UMAP manifold (Fig. S6D), suggesting that they are independent cell populations.

Seq-Scope also identified distinct cell types in the differentiated epithelial cell layer at the surface of the colonic mucosa (Fig. 6D-6F). The top layer of epithelial cells expressed an established marker for surface colonocytes, such as *Aqp8* (Fischer et al., 2001), Car4 (Borenshtein et al., 2009), Saa1 (Eckhardt et al., 2010) and *Lypd8* (Okumura et al., 2016) (Fig. 6G; Table S2), consistent with the former RNA *in situ* results (Borenshtein et al., 2009; Eckhardt et al., 2010; Fischer et al., 2001; Okumura et al., 2016). Some of the surface epithelial cells also expressed goblet cell-specific markers, such as *Zg16, Clca3, Fcgbp, Tff3* and *Spink3* (Haber et al., 2017; Pelaseyed et al., 2014) (Fig. 6G; Table S2). However, the distinction between the surface colonocytes and goblet cells was not so clear in the dataset processed with 10 μm square grids, because some colonocyte markers such as Lypd8 were abundantly found in the goblet grids (Fig. S6E). The colonocyte-goblet distinction became clearer when we used smaller grids, such as 7 μm square grids (Fig. S6F-S6I) and 5 μm square grids (Fig. S6J-S6M). Furthermore, when we directly plotted colonocyte and goblet cell markers onto the histological coordinate plane, the boundary between these two cell types was clearly visualized (Fig. 7C, S7D and S7E). These results indicate that acquiring high resolution is critical for precisely comprehending a single cell transcriptome from the ST dataset. In addition, Seq-Scope also identified hormone-secreting enteroendocrine cells (EEC; Fig. 6E) and identified their locations in the histological space (Fig. 6F, 7B and S7F).

Cell proliferative activities in the colon are higher in the crypt, where stem/dividing cells are located (Levine and Haggitt, 1989). Consistent with the notion, genes specifically expressed in proliferative cell cycle phases, such as S and G2/M phases (Nestorowa et al., 2016), were elevated in the crypt base area but relatively downregulated in the surface colonocyte area (Fig. 7D).

### Seq-Scope Identifies Colonic Cell Types in Subepithelial layers

Below the epithelial layer, there are connective tissue layers, including the lamina propria and submucosa, as well as external muscle layers. These layers also contain a diverse range of non-epithelial cell types (Fig. 6H). Seq-Scope also determined many additional cell types (Fig. 6I) and their spatial locations (Fig. 6J) in these layers. In addition to the smooth muscle cells described above, Seq-Scope identified fibroblasts (Fig. 7E and S7G) and enteric neurons (Fig. 7E and S7H). Seq-Scope also identified hematopoietically derived cells, such as myeloid cells (macrophages; Fig. S7I) and lymphoid cells (B-cells with different isotypes; Fig. S7J), which are located in the lamina propria (Fig. 6C, 6H, 6J and 7F). These different hematopoietic cell types are often in very close proximity (Fig. 7F, within μm distance), likely due to their functional interactions (Spencer and Sollid, 2016). However, since these cells expressed their unique cell type markers (Fig. 6K), Seq-Scope recognized them distinctively.

## Discussion

The data presented here demonstrate that Seq-Scope can visualize histological organization of the transcriptome architecture at multiple scales, including the tissue zonation level, cellular component level and even subcellular level. Equipped with an ultra-high-resolution output and an outstanding transcriptome capture output, Seq-Scope drew a clear boundary between different tissue zones, cell types and subcellular components. Previously existing technologies could not provide this level of clarity due to their low-resolution output and/or inefficiency in transcriptome capture.

Prior to Seq-Scope, HDST, which has a single barcode area of 9.3 μm^2^ (Vickovic et al., 2019), had been the technology that provided the highest resolution spatial transcriptomics (Asp et al., 2020; Liao et al., 2020; Zhou et al., 2020). However, HDST is seriously limited by its low transcriptome capture rate; a single barcoded area can catch only around 7 unique transcripts, even though the library was fully examined at around 90% saturation (Vickovic et al., 2019). In comparison, a single barcoded area of Seq-Scope, which is below 0.9 μm^2^ (less than 1/10 of HDST), can capture more unique transcripts at just around 70% (liver) and 42% (colon) saturation of library examination, leading to over 80 unique transcripts per 9.3 μm^2^ area. Therefore, Seq-Scope can produce 10 times more transcriptome output, in addition to providing 10 times more spatial resolution.

Compared to HDST, 10X VISIUM, currently the only commercialized ST solution, can provide much deeper information per barcode. Published data indicate that a single VISIUM barcode can identify approximately 5,000 (Liu et al., 2020) or 28,000 (Stickels et al., 2020) unique transcripts on average, depending on the tissue type. However, the barcode area of VISIUM is huge; it is a 55 μm-sided regular hexagon, occupying approximately 7800 μm^2^ of the histological area. Based on the current Seq-Scope output, it was estimated that approximately 78,000 transcripts could be identified in a single barcoded area of VISIUM. It is very difficult to perform a direct comparison because gastrointestinal tissues, examined in the current study, were not evaluated by these former technologies. Nevertheless, the big difference in the transcriptome output indicates that Seq-Scope has an outstanding transcriptome capture efficiency.

Current Seq-Scope colon data produced approximately 1,000 unique transcripts per 100 μm^2^ in the center-located tiles (tile ID: 2110-2114). Based on the current library sequencing saturation rate, we prospect that we may be able to obtain up to 2,000 unique transcripts per 100 μm^2^ area. This output is comparable to the most recent high-output ST technologies with 10 μm resolution, such as Slide-seqV2 (~550 per 100 μm^2^) (Stickels et al., 2020) and DBiT-seq (1,320-4,900 per 100 μm^2^) (Liu et al., 2020). Therefore, in addition to providing an unprecedented submicrometer resolution, Seq-Scope can reveal high-quality transcriptome information. Based on the high-resolution high-output performance, Seq-Scope was sufficient to perform single cell and subcellular studies of spatial transcriptome and reveal many biologically-interesting ST features from liver and colon slides.

While the current manuscript is in preparation, another technology, Stereo-Seq, claimed a nominal transcriptome resolution of 0.5-0.7 μm (Chen et al., 2021), which is similar to Seq-Scope. However, unlike Seq-Scope, Stereo-Seq suffered a low transcriptome capture efficiency, which is ~170 unique transcripts per 100 μm^2^ (less than 20% of current Seq-Scope output). Due to the low efficiency, most of the transcriptome studies in the Stereo-Seq work (Chen et al., 2021) were performed in 36 μm-sided square grids, which can contain more than 10 different single cells. Therefore, although Stereo-Seq was able to identify gross anatomical structures of embryonic organs and adult brain compartments (Chen et al., 2021), it did not reveal microscopic details such as single cell- or subcellular-level transcriptome information.

Several factors could have contributed to Seq-Scope’s high transcriptome capture efficiency. First, the dense and tight arrangement of barcoded clusters in Seq-Scope could have increased the transcriptome capture rate because they almost eliminated “blind spot” areas, which are substantial in the other ST methods using printed spots, beads or nanoballs. Second, unlike the bead-based ST methods, which produce a bumpy array surface, Seq-Scope produces a flat array surface, enabling direct interaction between the capture probe and tissue sample. Third, solid-phase amplification, limited by molecular crowding (Mercier and Slater, 2005), might have provided the two-dimensional concentration of RNA-capture probes ideal for the molecular interaction with tissue-derived RNA. Finally, biochemical strategies specific to our protocol, such as the secondary strand synthesis, retrieval, and amplification methods, could have increased the yield of transcriptome recovery.

Another benefit of Seq-Scope is its scalability and adaptability. Currently, we used the MiSeq platform for the HDMI-array generation; however, virtually any sequencing platforms using spatially localized amplification, such as Illumina platforms including GAIIx, HiSeq, NextSeq and NovaSeq, could be used for generation of the HDMI-array. The established technologies for DNA sequencing could be easily repurposed to provide high-resolution spatial barcoding. For instance, although MiSeq has fragmented imaging areas, each of which is limited to a 1 mm diameter circle, HiSeq2500 (Rapid-Run) and NovaSeq can provide approximately 90 mm^2^ and 800 mm^2^ of uninterrupted imaging area, respectively, which can offer a larger field of view. Newer sequencing methods are based on a patterned flow cell technology (Singer et al., 2019), which could provide a more defined spatial information for the HDMI-encoded clusters. However, it is also possible that patterned nanowells may limit the effective RNA capture area, leading to a lower RNA capture efficiency.

In terms of the cost, the current MiSeq-based HDMI-array can be generated at approximately $150 per mm^2^. The cost could be reduced further down to $11 per mm^2^ in HiSeq2500 or $2.6 per mm^2^ in NovaSeq, based on the current cost of sequencing. 30- and 40-nucleotide random seed sequence could provide a 1 quintillion and 1 septillion barcode diversities, respectively, which should be enough for spatially barcoding the wide imaging area surfaces. In terms of turnaround time, the HDMI-array generation takes less than a day after 1^st^-Seq, and library preparation could be completed within two days (three days in total except the sequencing). The procedure is straightforward and not laborious or technically demanding; correspondingly, a single researcher can simultaneously handle multiple samples. Therefore, Seq-Scope can make ultra-high-resolution ST accessible for any type and scale of basic science and clinical work.

Convenience in the data analysis pipeline is another strength of Seq-Scope. Most of the Seq-Scope analyses were seamlessly performed with widely-used standard software tools, such as Illumina MCS/RTA (Ravi et al., 2018), STARsolo (Dobin et al., 2013) and Seurat (Butler et al., 2018). High-resolution gene expression images produced by Seq-Scope could be handled like fluorescence microscope images, as we have demonstrated in this paper to visualize nuclear-cytoplasmic structure, liver zonation, colonic wall layers and interactions between single cell types and subtypes. Being relatively effortless in analytic perspective will be a hugely attractive factor for many experimental biologists.

The liver and colon datasets described in this paper are the first two datasets produced using the Seq-Scope technology. Having said that, there is a lot of room for further improvement. For instance, the MiSeq flow cell was not originally designed for ST (Ravi et al., 2018); therefore, exposing the cluster surface was physically challenging. Correspondingly, the cluster surface (HDMI-array) was substantially damaged during the initial flow cell disassembly, leading to many scratch-associated data losses in the liver dataset. When generating the colon dataset, we minimized the scratch-associated data losses by protecting the HDMI-array with a hydrogel filling method. In the future, designing a detachable flow cell for 1^st^-Seq could make this process even more straightforward.

The current HDMI-array is not encoded by the UMI sequence; therefore, we alternatively used random priming positions during the secondary strand synthesis as the alternative UMI. This strategy worked well for the transcriptome study. However, for detecting non-mRNA sequences, such as short micro RNA or antibody-attached oligonucleotides (Stoeckius et al., 2017), UMI-encoded HDMI-array could be more useful in the future. UMI-encoded HDMI-array could also enable tagmentation of the mRNA library, which could increase the diversity of the transcriptome read sequences (Hughes et al., 2020; Stickels et al., 2020). UMI encoding of the HDMI-array could be achieved by various methods, including oligonucleotide ligation (Vickovic et al., 2019) or template-based extension (Fig. S1F). We experimented on generating the UMI-encoded HDMI-array by template-based extension and found that both UMIs, produced by either random priming or array encoding, perform well in collapsing the PCR-multiplicated read sequences (Fig. S1G). In the future, the UMI-encoded HDMI-array could help apply Seq-Scope to multi-omics applications.

Finally, the data analysis pipeline could be further improved to fully utilize the high-resolution high-output characteristics of Seq-Scope. In the current study, a simple 10 μm square-gridding scheme was used to analyze our dataset through the standard cell type mapping (scRNA-seq) pipeline. Because the grid was smaller than the hepatocytes (25-30 μm in diameter) or colonocytes (10-15 μm in diameter), this analysis worked well and successfully identified most of the cell types present in the liver and colon. However, 10 μm square gridding is arbitrary, and a single grid could contain multiple cell types. Indeed, the hepatocyte markers, such as *Alb*, were found to be expressed in all grids examined in the dataset, although non-parenchymal cells are not supposed to express the hepatocyte-specific gene (Werner et al., 2015). The colon dataset also indicated that 10 μm square gridding was insufficient to separate Colonocyte and Goblet populations, which are located very close to each other in the histological space. This problem was ameliorated when we used smaller gridding or direct plotting onto the histology coordinate plane. However, smaller gridding also has a weakness because it can divide a single cell transcriptome into multiple grids, reducing transcriptome information available for each grid. An algorithm to detect the cell boundaries and precisely isolate single cell transcriptome could improve the Seq-Scope data analysis. Labeling cell boundaries through oligonucleotide-attached molecular probes, such as cell surface-targeting antibodies, could make this process more efficient and precise.

After further improvement in computational methods for single cell analysis, Seq-Scope has the potential to improve and complement current scRNA-seq approaches. scRNA-seq for solid tissues requires extensive tissue dissociation and single cell sorting procedures. These procedures create a very harsh condition for each cell type, which may alter the transcriptome with stress-associated responses. Furthermore, labile cell populations will be lost during tissue dissociation, and as a result, certain cell populations may be either over- or under-represented in the final dataset. Furthermore, there are many cell types, such as elongated myofibers and neurons, lipid-laden mature adipocytes, and cells tightly joined by the extracellular matrix and tight junctions, which are not amendable for conventional scRNA-seq analysis. By capturing the transcriptome directly from a frozen tissue slice, Seq-Scope can capture single cell transcriptome signatures from such difficult types of cells. Indeed, the Seq-Scope liver dataset revealed a couple of novel hepatocyte subpopulations undergoing tissue injury response, which were not formerly detected through scRNA-seq of normal and diseased liver tissues (Aizarani et al., 2019; Halpern et al., 2017; Park et al., 2020). This exemplifies the utility of Seq-Scope in identifying novel cell types from a solid tissue that were previously undetectable from traditional scRNA-seq; therefore, Seq-Scope also has the potential to complement and improve existing scRNA-seq technologies.

In sum, here we report on Seq-Scope technology, which enables transcriptome imaging at a microscopic resolution. A single run of Seq-Scope could produce microscopic gene expression imaging data for more than 20,000 genes. The vast amount of information produced by Seq-Scope would accelerate scientific discoveries and might lead to a new paradigm in molecular diagnosis.

## Author Contributions

C.S.C. performed experiments. J.X. performed computational analyses. S.R.P., J.E.H. and M.K. contributed to method development. G.J. and H.M.K. supervised J.X. in computational analyses. J.H.L. conceived the idea, led the project, and prepared the manuscript draft. All authors revised the manuscript and approved the final version.

## Acknowledgments

The authors thank Drs. Euisik Yoon, Hojoong Kwak, Chang H. Kim, Seungwon Jung and Yongsung Kim and Ms. Irene Hwang for their helpful advice and comments. We thank Psomagen Inc. and the University of Michigan Microbiome Core for their help in experiment optimization. The work was supported by the NIH (T32AG000114 to C.S.C., K01AG061236 to M.K, U01HL137182 to H.M.K. R01DK114131 and R01DK102850 to J.H.L., P30AG024824, P30AG013283, P30DK034933, P30DK089503, P30CA046592, P30AR069620, and U2CDK110768), the Chan Zuckerberg Initiative (to H.M.K.), the Frankel Cardiovascular Center Inaugural grant (to J.H.L. and M.K.), an American Association for the Study of Liver Diseases pilot research award (to J.H.L. and H.M.K.), Mcubed initiatives (to M.K., H.M.K. and J.H.L.), and Glenn Foundation grants (to J.H.L.).

## Experimental Procedures

### PART I. Experimental Implementation of Seq-Scope

#### Generation of Seed HDMI-oligo Library

Seq-Scope is initiated with the generation of a HDMI-oligo seed library (Fig. 1A and S1A). In the current report, we used two versions of the library – HDMI-DraI and HDMI32-DraI, whose sequences are provided below. The libraries have the same backbone structure with different lengths of HDMI sequences. HDMI is a sequence of random nucleotides designed to avoid the DraI digestion sites using Cutfree software (Storm and Jensen, 2018). HDMI32-DraI is an improved version of HDMI-DraI; however, for the liver and colon studies, HDMI-DraI was used. HDMI-DraI was generated by IDT as Ultramer oligonucleotides, while HDMI32-DraI was generated by Eurofins as Extremer oligonucleotides.

Backbone: (P5 sequence) (TR1: TruSeq Read 1) (HDMI) (HR1: HDMI Read 1) (Oligo-dT) (DraI) (DraI-adapter) (P7 sequence)

HDMI-DraI: CAAGCAGAAGACGGCATACGAGAT TCTTTCCCTACACGACGCTCTTCCGATCT NNVNNVNNVNNVNNVNNNNN TCTTGTGACTACAGCACCCTCGACTCTCGC TTTTTTTTTTTTTTTTTTTTTTTTTTT TTTAAA GACTTTCACCAGTCCATGAT GTGTAGATCTCGGTGGTCGCCGTATCATT

HDMI32-DraI: CAAGCAGAAGACGGCATACGAGAT TCTTTCCCTACACGACGCTCTTCCGATCT NNVNBVNNVNNVNNVNNVNNVNNVNNVNNNNN TCTTGTGACTACAGCACCCTCGACTCTCGC TTTTTTTTTTTTTTTTTTTTTTTTTTT TTTAAA GACTTTCACCAGTCCATGAT GTGTAGATCTCGGTGGTCGCCGTATCATT

#### HDMI-oligo Cluster Generation and Sequencing through MiSeq (1^st^-Seq)

HDMI-DraI or HDMI32-DraI was used as the ssDNA library and was sequenced in MiSeq using Read1-DraI as the custom Read1 primer. The Read1-DraI sequence is provided below.

Read1-DraI: ATCATGGACTGGTGAAAGTC TTTAAA AAAAAAAAAAAAAAAAAAAAAAAAAAA GCGAGAGTCGAGGGTGCTGTAGTCACAAGA

Read1-DraI has a complementary sequence covering HR1, Oligo-dT, DraI and DraI-adapter sequences of HDMI-DraI and HDMI32-DraI ssDNA libraries.

Initially, the libraries were sequenced using the MiSeq v2 nano platform to titrate the ssDNA library concentration to generate the largest possible number of confidently-sequenced HDMI clusters (Fig. S2A and S2B). After several rounds of optimization, HDMI-DraI was loaded at 100pM while HDMI32-DraI was loaded at 60-80pM. For actual implementation of Seq-Scope, the MiSeq v3 regular platform was used. MiSeq was performed in manual mode: 25bp single end reading (for HDMI-DraI) or 37bp single end reading (for HDMI32-DraI). The MiSeq runs were completed right after the first read without denaturation or re-synthesis steps. The flow cell was retrieved right after the completion of the single end reading steps. The MiSeq result was provided as a FASTQ file that has the HDMI sequence followed by the 5-base adapter sequence in TR1. The adapter sequence concordance was over 96% for all MiSeq results used in Seq-Scope. Thumbnail images of clusters, visualized using Illumina Sequencing Analysis Viewer, were used to inspect the cluster morphology and density (Fig. 2A, S2A and S2B).

The HDMI sequences contain 20-32 random nucleotides, which can produce 260 billion (20-mer in HDMI-DraI) or 1 quintillion (32-mer in HDMI32-DraI) different sequences. Due to this extreme diversity, the duplication rate of the HDMI sequence was extremely low (less than 0.1% of total HDMI sequencing results), even though the MiSeq identified more than 30 million HDMI clusters.

MiSeq has 38 rectangular imaging areas, which are called “tiles”. 19 tiles are on the top of the flow cell, while the other 19 tiles are on the bottom of the flow cell (Fig. S2C; tiles 2101-2119). For each sequencing output, the tile number and XY coordinates of the cluster from which the sequence originates can be found in the FASTQ output file of MiSeq. Only the bottom tiles were used for Seq-Scope analysis because the top tiles were destroyed during the flow cell disassembly.

#### Processing MiSeq Flow Cell into the HDMI-array

After 1^st^-Seq, the MiSeq flow cell was further processed to convert HDMI-containing clusters to HDMI-array (Fig. 1D). The flow cell retrieved from the MiSeq run was washed with nuclease-free water 3 times. Then, the flow cell was treated with DraI enzyme cocktail (1U DraI enzyme (#R0129, NEB) in 1X CutSmart buffer) in 37 °C overnight to completely cut out the P5 sequence and expose oligo-dT. The flow cell was then loaded with exonuclease I cocktail (1U Exo I enzyme (#M2903, NEB) in 1X Exo I buffer) in 37 °C for 45 min to eliminate P5 primer lawn and other non-specific ssDNA. P7-bound HDMI-DraI oligonucleotides make a duplex with Read1-DraI, so they were protected from Exo I digestion. Then, the flow cell was washed with water 3 times, 0.1N NaOH 3 times (each with 5 min incubation at room temperature, to denature and eliminate the Read1-DraI primer), 0.1M Tris (pH7.5, to neutralize the flow cell channel) 3 times (each with brief wash), and then water 3 times (each with a brief wash).

#### HDMI-array Disassembly

Then, the flow cell was disassembled so that the HDMI-array was exposed to the outside and could be attached to tissue sections. To protect the HDMI-array, agarose hydrogel (BP160, Fisher) was used to fill the flow cell channel before disassembly (for the colon dataset). 1.5% agarose suspension was prepared in water and incubated at 95 °C for 1 min. The resulting 1.5% melted agarose solution was loaded into the flow cell and chilled to solidify the gel. Using the Tungsten Carbide Tip Scriber (IMT-8806, IMT), the top glass layer was destroyed to expose the bottom layer for tissue attachment. Agarose hydrogel filling was removed by extensive washing with warm distilled water. The top-exposed flow cell, HDMI-array, was then ready for tissue attachment.

#### Tissue Samples

The liver and colon samples were from our recent studies (Cho et al., 2019; Ro et al., 2016). The livers were collected from 8 week-old control (*Depdc5*^*F/F*^/*Tsc1*^*F/F*^, male) and *TD* (Alb-Cre/*Depdc5*^*F/F*^/*Tsc1*^*F/F*^, female) mice (Cho et al., 2019). The colons are from 8-week-old C57BL/6 wild-type male mice (Ro et al., 2016).

#### Tissue Sectioning, Attachment and Fixation

OCT-mounted fresh frozen tissue was sectioned in a cryostat (Leica CM3050S, −20 C) at a 5° cutting angle and 10 μm thickness. The tissues were maneuvered onto the HDMI-array from the cutting stage (Fig. 1E). The tissue-HDMI-array sandwich was moved to room temperature, and the tissues were fixed in 4% formaldehyde (100 μl, diluted from the EM-grade 16% paraformaldehyde (#15170, Electron Microscopy Sciences)) for 10 min.

#### Tissue Imaging and mRNA release

The tissues were incubated 2 min in 100 μl isopropanol, and then stained with 80 μl hematoxylin (S3309, Agilent) for 5 min. After washing with water, the tissues were treated with 80 μl bluing buffer (CS702, Agilent) for 2 min. After washing with water, the tissues were treated with buffered eosin (1:9 = eosin (HT110216, Sigma): 0.45M Tris-Acetic buffer (pH 6.0)). After washing with water, the tissues were dried and mounted in 85% glycerol. The tissues were then imaged under a light microscope (MT6300, Meiji Techno). To release RNAs from the fixed tissues, the tissues were treated with 0.2U/uL collagenase I (17018-029, Thermo Fisher) at 37 °C 20 min, and then with 1mg/mL pepsin (P7000, Sigma) in 0.1M HCl at 37 °C for 10 min, as previously described (Salmen et al., 2018).

#### Reverse Transcription

The tissue was washed with 40 μl 1X RT buffer containing 8 μl Maxima 5x RT Buffer (EP0751, Thermofisher), 1 μl RNase Inhibitor (30281, Lucigen) and 31 μl water. Subsequently, reverse transcription (Fig. 1F and S1B) was performed by incubating the tissue-attached HDMI-array in 40 μl RT reaction solution containing 8 μl Maxima 5x RT Buffer (EP0751, Thermofisher), 8 μl 20% Ficoll PM-400 (F4375-10G, Sigma), 4 μl 10mM dNTPs (N0477L, NEB), 1 μl RNase Inhibitor (30281, Lucigen), 2 μl Maxima H- RTase (EP0751, Thermofisher), 4 μl Actinomycin D (500ng/μl, A1410, Sigma-Aldrich) and 13 μl water. The RT reaction solution was incubated at 42 °C overnight.

#### Tissue Digestion

The next day, the RT solution was removed, and the tissue was submerged in the exonuclease I cocktail (1U Exo I enzyme (#M2903, NEB) in 1X Exo I buffer) and incubated at 37 °C for 45 min to eliminate DNA that did not hybridize with mRNA. Then the cocktail was removed and the tissues were submerged in 1x tissue digestion buffer (100 mM Tris pH 8.0, 100 mM NaCl, 2% SDS, 5 mM EDTA, 16 U/mL Proteinase K (P8107S, NEB). The tissues were incubated at 37 °C for 40 min.

#### Secondary Strand Synthesis and Purification

After the tissue digestion, the HDMI-array was washed with water 3 times, 0.1N NaOH 3 times (each with 5 min incubation at room temperature), 0.1M Tris (pH7.5) 3 times (each with a brief wash), and then water 3 times (each with a brief wash). This step eliminated all mRNA from the HDMI-array.

After washing steps, The HDMI-array was treated with a secondary strand synthesis mix (18 μl water, 3 μl NEBuffer-2, 3 μl 100 μM Truseq Read2-conjugated Random Primer with TCA GAC GTG TGC TCT TCC GAT CTN NNN NNN NN sequence (IDT), 3 μl 10 mM dNTP mix (N0477, NEB), and 3 μl Klenow Fragment (exonuclease-deficient; M0212, NEB). The HDMI-array was incubated at 37 °C for 2 hr in a humidity-controlled chamber.

After secondary strand synthesis (Fig. 1G), the HDMI-array was washed with water 3 times to remove all DNAs that were not bound to the HDMI-array, so that each HDMI molecule corresponded to each single copy of the secondary strand. Then the HDMI-array was treated with 30 μl 0.1 N NaOH to elute the secondary strand. The elution step was duplicated to collect 60 μl (in total) of the secondary strand product. The 60 μl secondary strand product was neutralized by mixing with 30 μl 3 M potassium acetate, pH5.5.

The volume of the neutralized secondary strand product was adjusted to 100 μl by adding ~10 μl water. The solution was then subjected to AMPure XP purification (A63881, Beckman Coulter) using a 1.8X bead/sample ratio, according to the manufacturer’s instruction. The final elution was performed using 40 μl water.

#### Library Construction and Sequencing (2^nd^-Seq)

First-round library PCR was performed using Kapa HiFi Hotstart Readymix (KK2602, KAPA Biosystems) in a 100 μl reaction volume with 40 μl secondary strand product as the template and forward (TCT TTC CCT ACA CGA CGC*T*C) and reverse (TCA GAC GTG TGC TCT TCC*G*A) primers at 2 μM. PCR condition: 95 °C 3 min, 13-15 cycles of (95 °C 30 sec, 60 °C 1 min, 72 °C 1 min), 72 °C 2 min and 4 °C infinite. PCR products were purified using AMPure XP in a 1.2X bead/sample ratio.

Second-round library PCR (Fig. 1H) was performed using Kapa HiFi Hotstart Readymix (KK2602, KAPA Biosystems) in 100 μl reaction volume with 10 μl of 2 nM first-round PCR product as a template and forward (AAT GAT ACG GCG ACC ACC GAG ATC TAC ACT CTT TCC CTA CAC GAC GCT CT*T *C) and reverse (CAA GCA GAA GAC GGC ATA CGA GAT [8-mer index sequence] GTG ACT GGA GTT CAG ACG TGT GCT CTT CC*G *A) primers at 1 μM. PCR condition: 95 °C 3 min, 8-9 cycles of (95 °C 30 sec, 60 °C 30 sec, 72 °C 30 sec), 72 °C 2 min and 4 °C infinite. PCR products were purified using agarose gel elution for all products between 400-850bp size, using the Zymoclean Gel DNA Recovery Kit (D4001, Zymo Research) according to the manufacturer’s recommendation. Then the elution products were further purified using AMPure XP in a 0.6X-0.7X bead/sample ratio. The pooled libraries were subjected to paired-end (100-150bp) sequencing in the Illumina and BGI platforms at AdmeraHealth Inc., Psomagen Inc., and Beijing Genome Institute. The HDMI discovery plot assessments indicated that all sequencing platforms worked well for analyzing Seq-Scope data.

#### cDNA Labeling Assay

To label cDNAs on the HDMI-array, all the steps were identically performed as described above, except that, after mRNA release, the HDMI array was subjected to cDNA labeling assay (Salmen et al., 2018) instead of library generation procedures. After mRNA release, the tissue-attached HDMI array was incubated in 40uL fluorescent reverse transcription solution containing 13 μl water, 8 μl Maxima 5x RT Buffer (EP0751, Thermofisher), 8 μl 20% Ficoll PM-400 (F4375-10G, Sigma), 0.8 μl 100mM dATP (from 0446S, NEB), 0.8 μl 100mM dTTP (from 0446S, NEB), 0.8 μl 100mM dGTP (from 0446S, NEB), 0.1 μl 100mM dCTP (from 0446S, NEB), 1.5 μl 6.45mM Cy3-dCTP (B8159, APExBIO), 1 μl RNase Inhibitor (30281, Lucigen), 4 μl Actinomycin D (500ng/μl, A1410, Sigma-Aldrich) and 2 μl Maxima H- RTase (EP0751, Thermofisher). Reverse transcription was performed at 42°C overnight.

Then, the cocktail was removed and the tissues were submerged in 1x tissue digestion buffer (100 mM Tris pH 8.0, 100 mM NaCl, 2% SDS, 5 mM EDTA, 16 U/mL Proteinase K (P8107S, NEB)). The tissues were incubated at 37 °C for 40 min. After washing the HDMI-array surface with water 3 times, it was mounted in 80% glycerol and then observed under a fluorescent microscope (Meiji).

#### Generation and Testing of UMI-encoded HDMI-array

UMI-encoded HDMI array was generated using the HDMI-TruEcoRI library, which is similar to the other ssDNA libraries described above but it does not have an oligo-dT sequence (Fig. S1F).

Backbone: (P5 sequence) (TR1: TruSeq Read 1) (HDMI) (HR1B: HDMI Read 1B) (EcoRI) (EcoRI adapter) (P7 sequence)

HDMI-TruEcoRI: CAAGCAGAAGACGGCATACGAGAT TCTTTCCCTACACGACGCTCTTCCGATCT HNNBNBNBNBNBNBNBNNNN CCCGTTCGCAACATGTCTGGCGTCATA GAATTC CGCAGTCCAG GTGTAGATCTCGGTGGTCGCCGTATCATT

For MiSeq running, Read1-EcoRI was used as the read 1 primer.

Backbone: (EcoRI adapter) (EcoRI) (HR1B)

Read1-EcoRI: CTGGACTGCG GAATTC TATGACGCCAGACATGTTGCGAACGGG

The library was sequenced using MiSeq v2 nano platform at 100pM concentration and generated 1.4 million sequenced HDMI clusters per mm^2^. MiSeq was performed in manual mode, 25bp single end reading, using the Read1-EcoRI as the custom Read 1 primer. The flow cell was retrieved right after the completion of the single end reading step. Then, the MiSeq flow cell was processed to attach UMI and oligo-dT sequences to the HDMI clusters. The flow cell was washed with water 3 times and then loaded with EcoRI-HF cocktail (1U EcoRI-HF (R3101, NEB) in 1X CutSmart NEB buffer) to cut out the P5 sequence. After 37 °C overnight incubation, the flow cell was washed with water 3 times, 0.1N NaOH 3 times (each with 5 min incubation at room temperature), 0.1M Tris (pH 7.5) 3 times, and then water 3 times. The flow cell was then loaded with 1X Phusion Hot Start II High-Fidelity Mastermix (F565S) containing 5 μM of UMI-oligo (sequence provided below).

Backbone: (oligo-dA) (UMI) C (HR1B)

UMI-Oligo: AAAAAAAAAAAAAAAAAAAAAAAAAAAAAA NNNNNNNN C TATGACGCCAGACATGTTGCGAACGGG

The flow cell was then incubated at 95 °C for 5 min, 60 °C for 1 min and 72 °C for 5 min. Then, the flow cell was loaded with an exonuclease I cocktail (see above for composition) and incubated for 45 min at 37 °C. The flow cell was then washed with water 3 times, 0.1N NaOH 3 times (each with 5 min incubation at room temperature), 0.1M Tris (pH 7.5) 3 times, and then water 3 times. This completes the generation of the UMI-encoded HDMI-array.

Performance of the UMI-encoded HDMI-array was tested using 2 μg total RNA purified from mouse liver, using the same reverse transcription and library preparation method described above (but without the tissue slice). The library was sequenced in Illumina HiSeqX and HiSeq4000 platforms.

### PART II. Computational Analysis of Seq-Scope data

#### Input Data

There are three experimental outputs from Seq-Scope, which will serve as input data for downstream computational analysis. (1) HDMI sequence, tile and spatial coordinate information from 1^st^-Seq, (2) HDMI sequence, coupled with cDNA sequence from 2^nd^-Seq, and (3) Histological image obtained from H&E staining of the tissue slice.

#### Tissue Boundary Estimation

To estimate the tissue boundary, the HiSeq data were joined into MiSeq data according to their HDMI sequence. As a result, for each of the HiSeq data whose HDMI was found from MiSeq, the tile number and XY coordinates were assigned. Finally, using a custom python code, an HDMI discovery plot was generated to visualize the density of HiSeq HDMI in a given XY space of each tile (Fig. S1C). The density plots were manually assigned to the corresponding H&E images (Fig. 2C, S2D and S2E).

#### Read Alignment and Generation of Digital Gene Expression Matrix

Read alignment was performed using STAR/STARsolo 2.7.5c (Dobin et al., 2013), from which the digital gene expression (DGE) matrix was generated. From MiSeq data, HDMI sequences of clusters located on the bottom tile were extracted and used as a “whitelist” for the cell (HDMI) barcode after reverse complement conversion. The first 20 (HDMI-DraI version) or 30 (HDMI32-DraI) basepairs of HiSeq data Read 1 were considered as the cell (HDMI) barcode.

Due to the extensive washing steps after secondary strand synthesis, it was expected that each single molecule of HDMI-cDNA hybrid would lead to one secondary strand in the library. Therefore, the first 9-mer of Read 2 sequence, which is derived from the Randomer sequence, could serve as a proxy of the unique molecular identifier (UMI). Therefore, the first 9 basepairs of HiSeq Read 2 data were copied to Read 1 and used as the unique molecular identifier (UMI). Read 2 was trimmed at the 3’end to remove polyA tails of length 10 or greater and was then aligned to the mouse genome (mm10) using the Genefull option with no length threshold and no cell filtering (Fig. S1D). For the genes whose expression couldn’t be monitored by the Genefull option, the Gene option was used to generate the gene expression discovery plots.

For saturation analysis, multiple read alignments were performed using various subsets of the 2^nd^-Seq results. The alignment output values were plotted in a graph (Fig. S2P) to generate a saturation curve in Graphpad Prism 8 (Graphpad Software, Inc.). Hyperbolic regression was used to estimate the total unique transcript number in the liver (59,799,349 to 63,511,208; 95% confidence interval) and colon (107,954,654 to 123,477,902; 95% confidence interval) Seq-Scope libraries.

#### Analysis of Spliced and Unspliced Gene Expression

To obtain separate read counts for spliced and unspliced transcripts, we used the Velocyto (La Manno et al., 2018) option in the STARsolo software (Fig. S1E). All spliced or unspliced mRNA reads were plotted onto the histological coordinate plane to identify nuclear-cytoplasmic structure (see below in “Visualization of Spatial Gene Expression). To test the reproducibility of the image analysis, all genes were randomly divided into three groups, and spliced and unspliced read counts were obtained independently. Images were compared with each other to calculate Pearson’s correlation coefficients in NIH ImageJ using Just Another Colocalization Plugin (JACoP) (Bolte and Cordelieres, 2006). Abundances of nuclear-specific (*Malat1, Neat1* and *Mlxipl*) and mitochondrial-encoded (all genes whose name start with “mt-“) transcripts were also analyzed using the same method. The correlation coefficients were assembled and presented in a heat map produced by Graphpad Prism 8 (Graphpad Software Inc.).

#### Data Binning and Clustering Analysis

Data binning was performed by dividing the imaging space into 100 μm^2^ (10 μm-sided) square grids and collapsing all HDMI-UMI information into one barcode per grid. Alternatively, data binning was also performed using 49 μm^2^ (7 μm-sided) square grids or 25 μm^2^ (5 μm-sided) square grids. The binned DGE matrix was analyzed in the Seurat v4 package (Butler et al., 2018). Feature number threshold was applied to remove the grids that corresponded to the area that was not overlaid by the tissue or was extensively damaged through scratches. Data were normalized using regularized negative binomial regression implemented in Seurat’s SCTransform function. Clustering was performed using the shared nearest neighbor modularity optimization implemented in Seurat’s FindClusters function, using resolutions between 0.6 and 1.2. Clusters with apparently mixed cell types (cluster 8 in Fig. S5A, cluster 6 and 8 in Fig. S6B) were subjected to an additional round of clustering. UMAP (Becht et al., 2019) manifold, also built in the Seurat package, was used to assess clustering performance. Top 50 markers from each cluster, identified through the FindAllMarkers function, were used to identify cell types that existed within the grid area. Then the clusters were visualized in the UMAP manifold or the histological space using DimPlot and SpatialDimPlot functions, respectively. Raw and normalized transcript abundance in each tile, cluster and spatial grid was visualized through the VlnPlot, DotPlot, FeaturePlot and SpatialFeaturePlot functions built in the Seurat package.

#### Visualization of Spatial Gene Expression

Spatial gene expression was visualized using a custom python code. Raw digital expression data of the queried gene (or gene list) were plotted onto the coordinate plane according to their HDMI spatial index. Gene expression densities were plotted as a ~3 μm-radius circle at a transparency alpha level of between 0.005 and 0.25. In spatial gene expression images with the white background, the intensity of the colored spot indicates the abundance of transcripts around the spot location. Spatial gene expression images with the black background were created for genes or gene lists of high expression values to make it easy to adjust the linear range of gene expression density and to overlay gene expression densities of different queries with different pseudo-color encoding. The inverse image of the greyscale plot was pseudo-colored with red, blue or green, and the image contrast was linearly adjusted to highlight the biologically relevant spatial features. Finally, different pseudo-colored images were overlaid together to compare the gene expression patterns in the same histological coordinate plane. Cell cycle-specific genes, such as S phase- and G2/M phase-specific gene lists (Nestorowa et al., 2016), were retrieved from the Seurat package, and their mouse homologues were identified using the biomaRt package (Durinck et al., 2009).

#### UMI Efficiency Test

Efficiencies of UMI-encoding methods for collapsing duplicate read counts were evaluated using the data produced from the “Generation and Testing of UMI-encoded HDMI-array” section. UMI encoded by the HDMI-array (UMI_Array; 49^th^-57^th^ positions of Read 1) and UMI encoded by the Random primed position (UMI_Randomer; 1^st^-9^th^ positions of Read 2) were identified from the 2^nd^-Seq results. Uncollapsed read count, read count collapsed with UMI_Array, and read count collapsed with UMI_Randomer were calculated for all the HDMI sequences observed, and their relative abundances were presented in a line graph (Fig. S1G). The result indicates that both UMI_Array and UMI_Randomer are efficient in collapsing duplicate read counts of 2^nd^-Seq results.

## Supplemental Figure Legends

**Fig S1.**
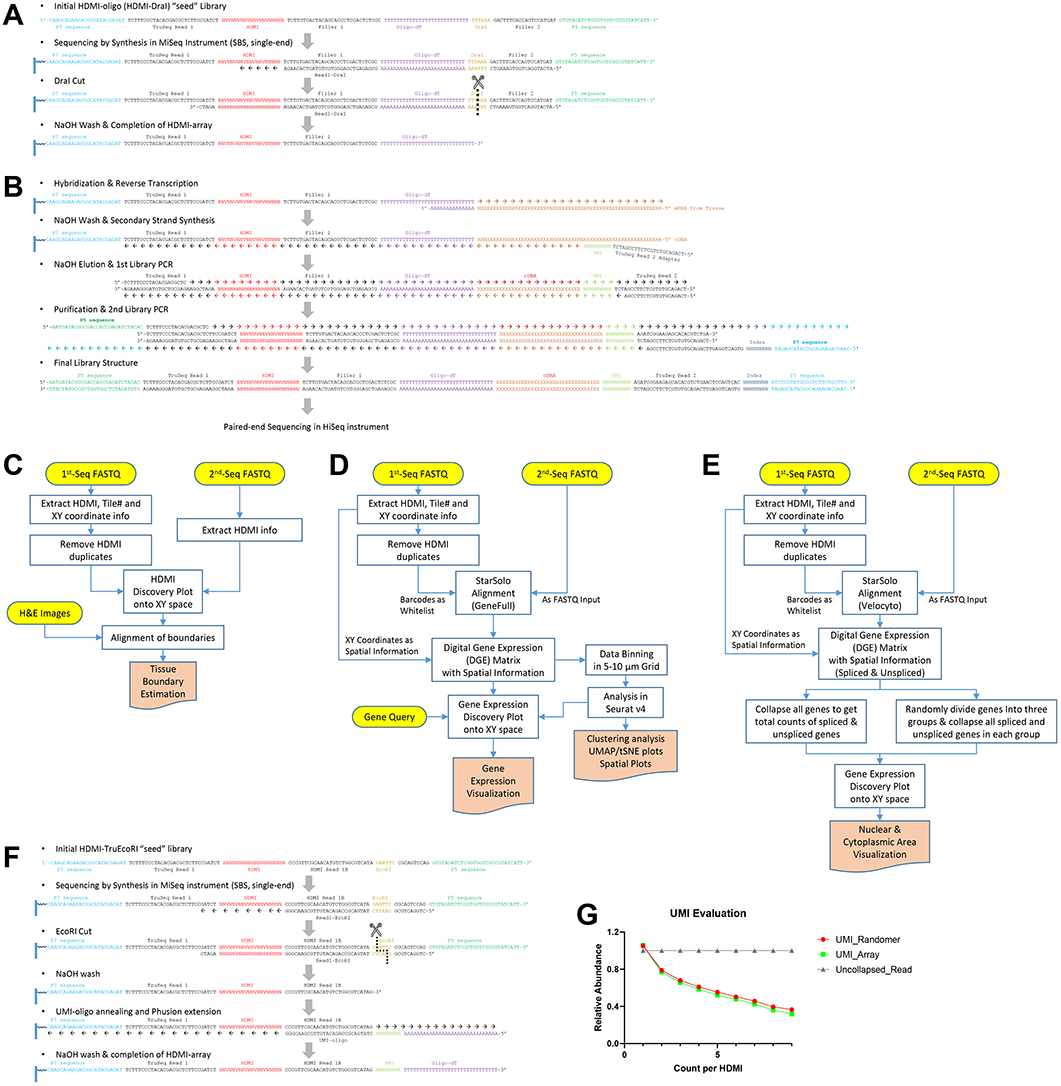
Seq-Scope Workflow. (A and B) Chemistry workflow for generating HDMI-array in 1^st^-Seq (A), and using the HDMI-array for constructing library for 2^nd^-Seq (B). The 2^nd^-Seq library is subjected to the standard next-generation sequencing workflow in Illumina and BGI platforms. See Experimental Procedures for details. (C-E) Bioinformatics workflow for estimating tissue boundaries (C), visualizing and analyzing spatial gene expression patterns (D), and determining nuclear and cytoplasmic areas (E). See Experimental Procedures for details. (F) Chemistry workflow for generating UMI-encoded HDMI-array in 1^st^-Seq. (G) Evaluation of UMI-encoding methods based on either random priming (UMI_Randomer) or array encoding (UMI_Array). The number of HDMI with multiple read counts was efficiently reduced by either UMI_Randomer- or UMI_Array-based collapsing methods.

**Fig S2.**
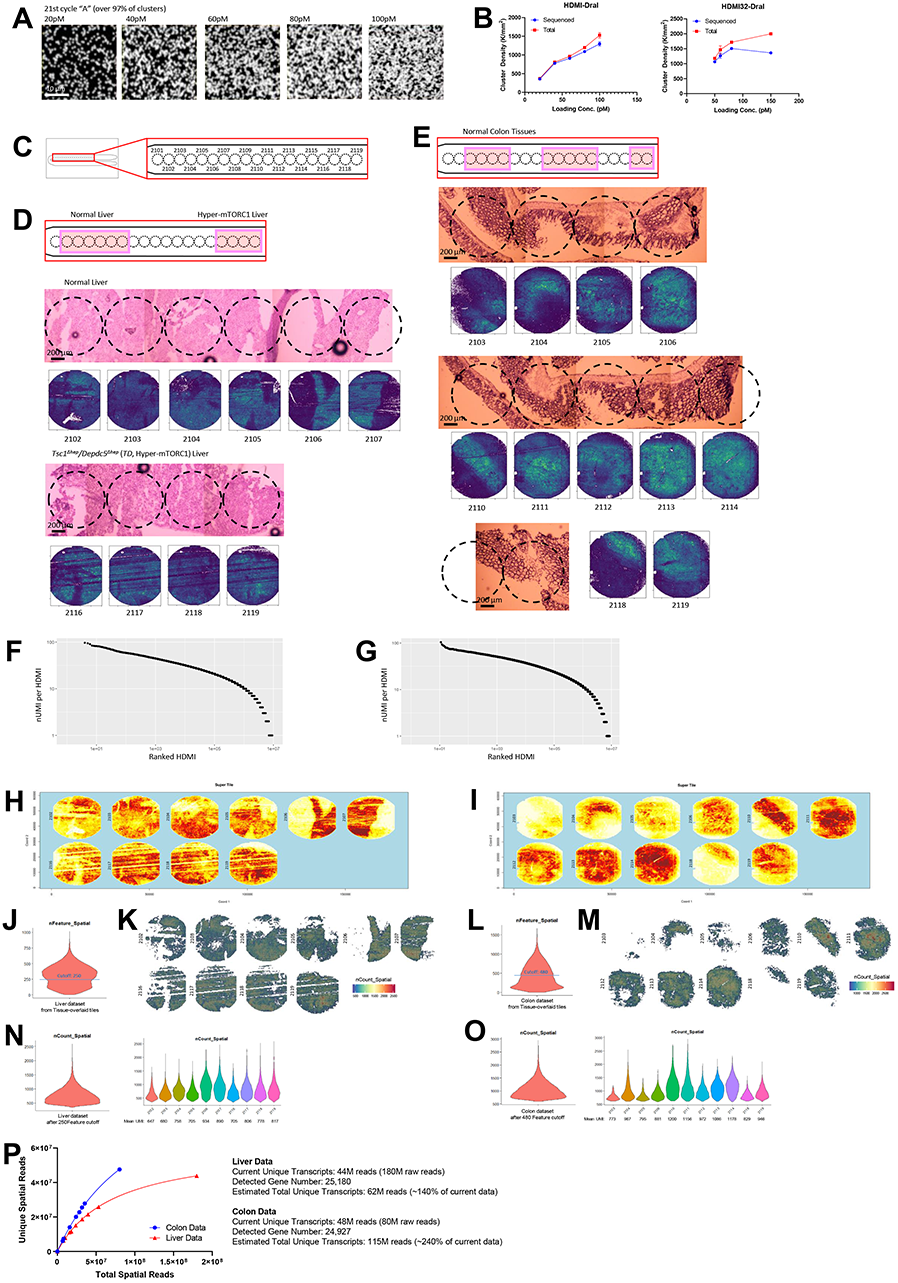
Seq-Scope Performance. (A) Representative images of HDMI clusters in the HDMI-array, retrieved from the Illumina sequence analysis viewer (SAV). Each picture visualizes “A” intensity at the 21^st^ cycle of the 1^st^-Seq SBS, where over 97% of HDMI clusters exhibit fluorescence. (B) Titration of HDMI-oligo library loading concentration for obtaining maximum number of sequenced clusters. Total (red) and sequenced (blue) cluster numbers were presented for indicated 1^st^-Seq conditions. Data are presented as mean ± SEM. (C) Schematic diagram depicting tile arrangement in bottom surface of MiSeq v3 regular flow cell. (D and E) Schematic diagram visualizes the tiles which were attached to the indicated liver (D, top) or colon (E, top) tissues. On the bottom, H&E staining images (upper) and their corresponding HDMI discovery plots (lower) were presented. (F and G) Knee plots depicting the distribution of all HDMI discovered from 2^nd^-Seq and the number of unique transcripts (nUMI) discovered per each HDMI molecule. Outlier points (n≤10 each) with more than 100 nUMI were not plotted in the graph area. Both liver (F) and colon (G) datasets were analyzed. (H and I) Spatial density plots of the gridded dataset depicting the number of UMIs discovered from indicated 10 μm square grids. (J-O) Violin plots depicting the distribution of the number of gene feature (nFeature) across the 10 μm square grids (J and L). Setting a 250 (liver) or 480 (colon) cutoff for these tiles isolated grid units covered by the tissue area (K and M), each of which contains up to 1,200 UMIs (N and O). Both liver (J, K and N) and colon (L, M and O) datasets were analyzed. (P) Saturation analysis of liver (red) and colon (blue) Seq-Scope dataset.

**Fig S3.**
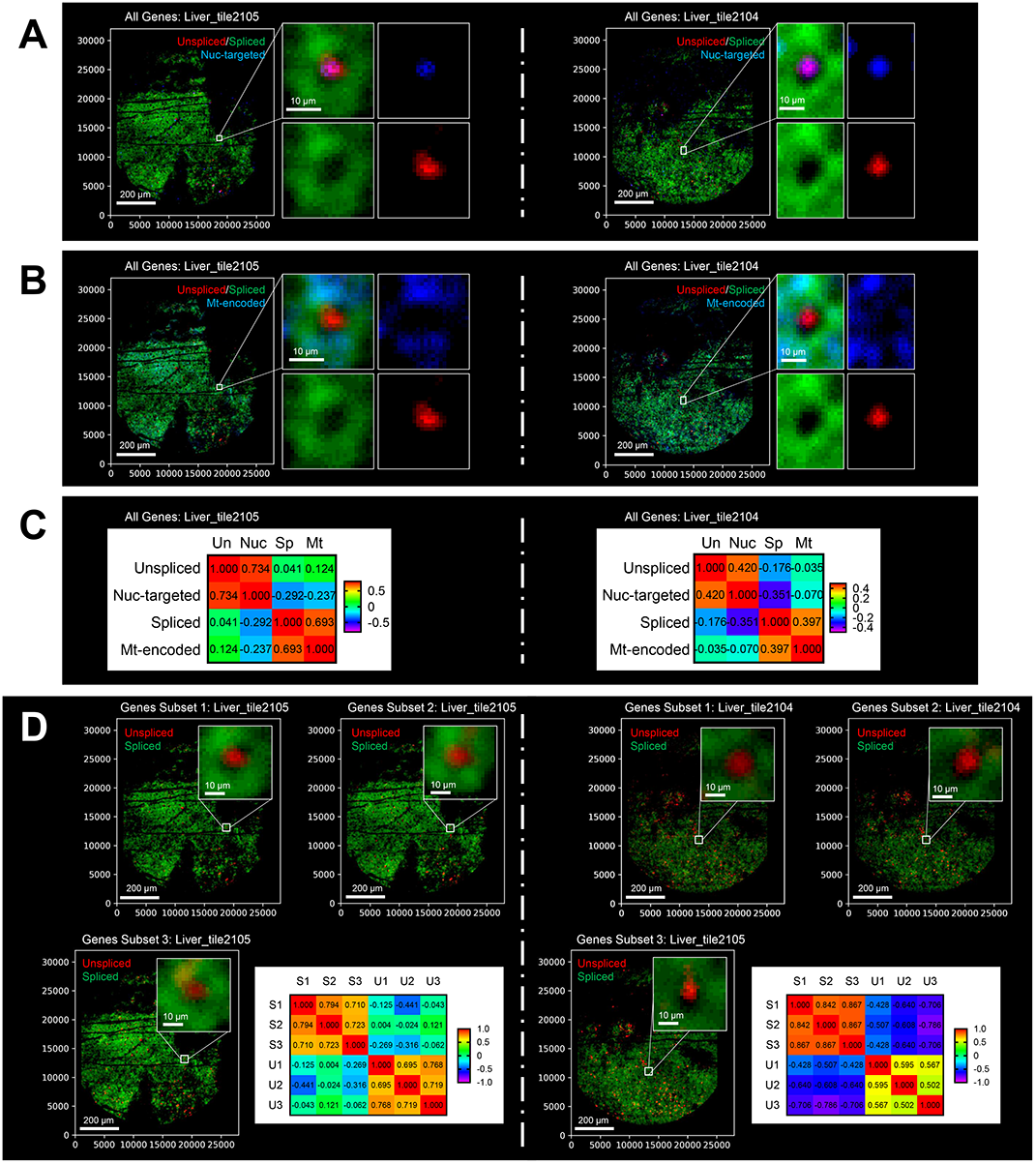
Seq-Scope Visualizes Nuclear/Mitochondrial/Cytoplasmic Subcellular Architecture. This figure provides additional examples of Seq-Scope output that visualizes the nuclear/mitochondrial/cytoplasmic subcellular architecture (A) Spatial plot of all unspliced and spliced transcripts, as well as RNA species that are known to localize to nucleus in liver tissue (Nuc-targeted; *Malat1, Neat1 and Mlxipl*). (B) Spatial plot of all unspliced and spliced transcripts, as well as RNA species that are encoded by mitochondrial genome (Mt-encoded). (C) Pearson correlations (*r*) between the indicated transcript intensities in the single cell area were presented as a heat map. (D) Spatial plot of unspliced and spliced transcript in three independent subsets of genes (Gene Subset 1-3). Pearson correlations (*r*) between these transcript intensities were presented as a heat map. S1-3, Spliced 1-3; U1-3, Unspliced 1-3.

**Fig S4.**
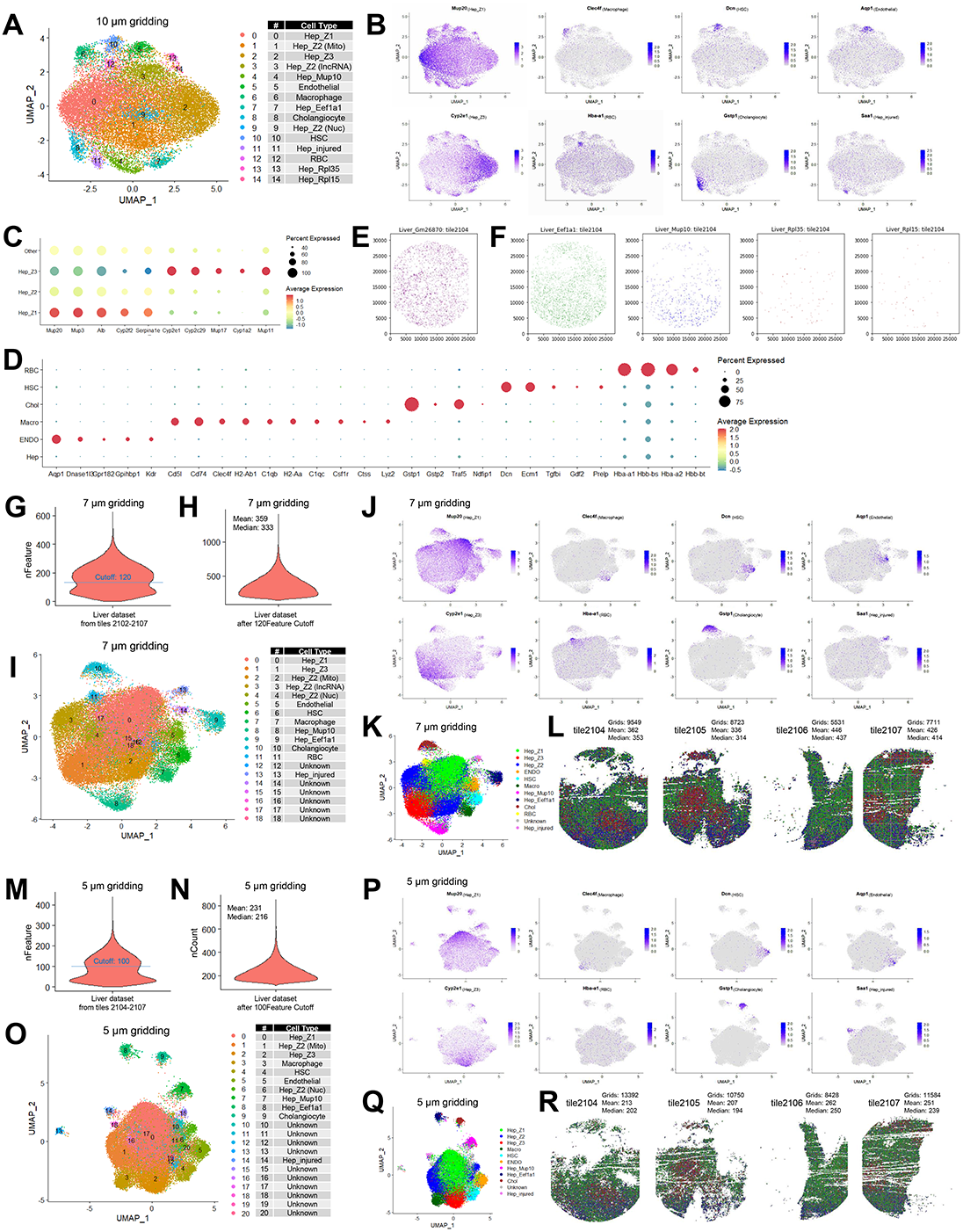
Seq-Scope Identifies Diverse Cell Types and Subtypes Present in Normal Liver. (A-D) From the normal liver dataset binned with 10 μm-sided square grids, a UMAP plot visualizing all clusters (A), UMAP plots visualizing expression of indicated genes across the grids (B), and dot plots visualizing cluster-specific expression of liver zonation (C) and cell type (D) markers are presented. (E and F) Spatial plot of indicated transcripts on histological coordinate plane. (G-J) Number of gene features (G, nFeatures) and UMI counts (H, nCounts; after nFeatures cutoff at 120) were calculated across the indicated tiles of liver Seq-Scope dataset, binned using 7 μm square grids. From this dataset, a UMAP plot visualizing all clusters (I), UMAP plots visualizing expression of indicated genes across the grids (J), a UMAP plot visualizing cell type-assigned clusters (K) and its associated spatial plots (L) are presented. Grid numbers, as well as mean and median UMI counts per grid unit, were provided (L). (M-R) Number of gene features (M, nFeatures) and UMI counts (N, nCounts; after nFeatures cutoff at 100) were calculated across the indicated tiles of liver Seq-Scope dataset, binned using 5 μm square grids. From this dataset, a UMAP plot visualizing all clusters (O), UMAP plots visualizing expression of indicated genes across the grids (P), a UMAP plot visualizing cell type-assigned clusters (Q) and its associated spatial plots (R) are presented. Grid numbers, as well as mean and median UMI counts per grid unit, were provided (R).

**Fig S5.**
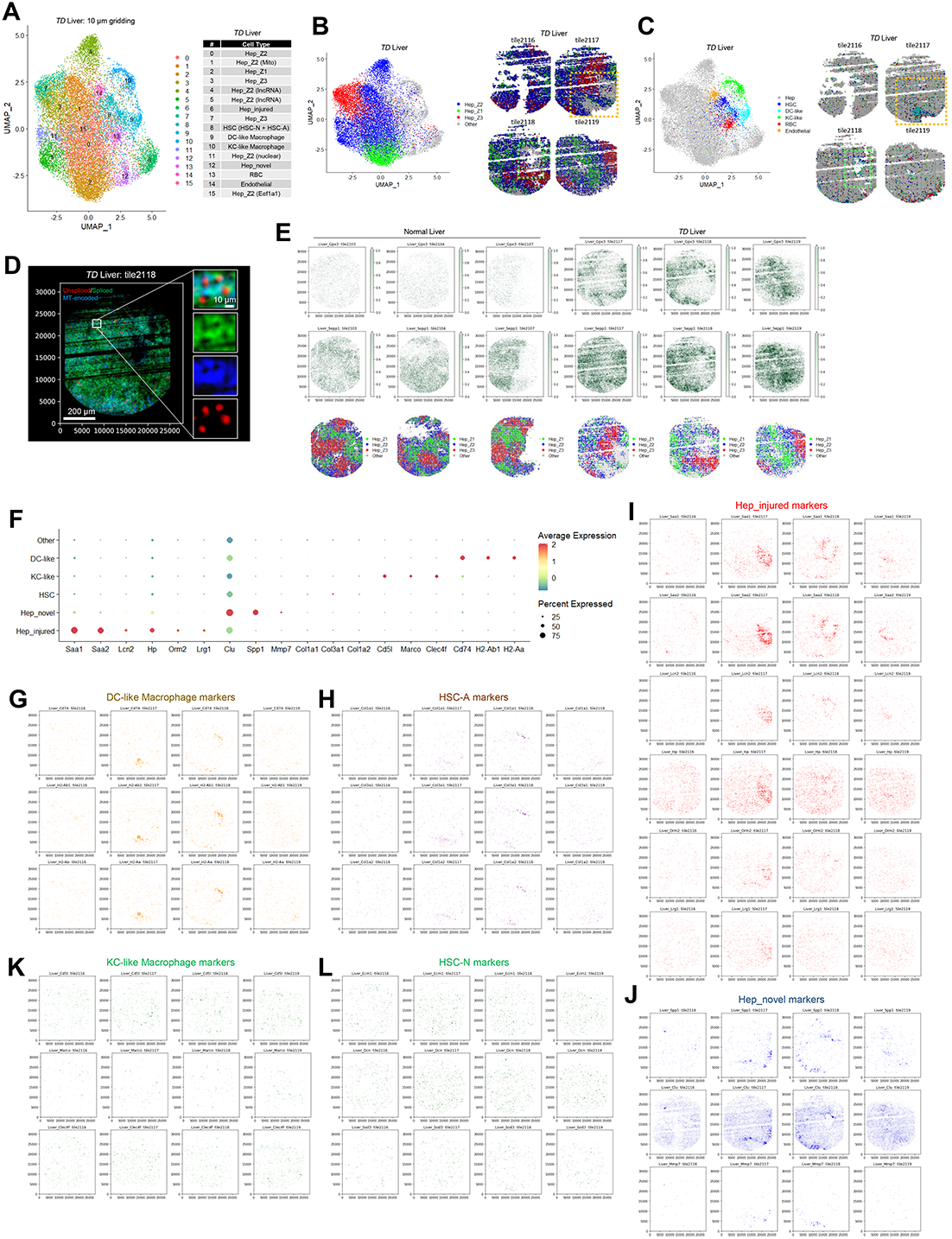
Seq-Scope Analysis of Liver Injury and Inflammation. (A-C) From the *TD* liver dataset binned with 10 μm-sided square grids, UMAP plots visualizing all clusters (A), zonated hepatocytes (B) and non-parenchymal cells (C) are presented. (D) Spatial plots of unspliced, spliced and mitochondrial transcripts visualize subcellular structures. (E) Expression of oxidative stress-responsive genes, *Gpx3* and *Sepp1*, was examined in normal and *TD* liver using spatial plotting. Hepatocyte zonation is plotted in the bottom panel as a reference. *Gpx3* and *Sepp1* were specifically induced in zone 1 hepatocytes of *TD* liver. (F) Dot plot depicting cluster-specific expression of cell type marker genes. (G-J) Spatial plots visualizing expression of marker genes for dendritic cell-like macrophages (G), activated HSCs (H), hepatocytes showing injury response (I), hepatocytes showing novel injury response (J), Kupffer cell-like macrophages (K) and normal HSCs (L).

**Fig S6.**
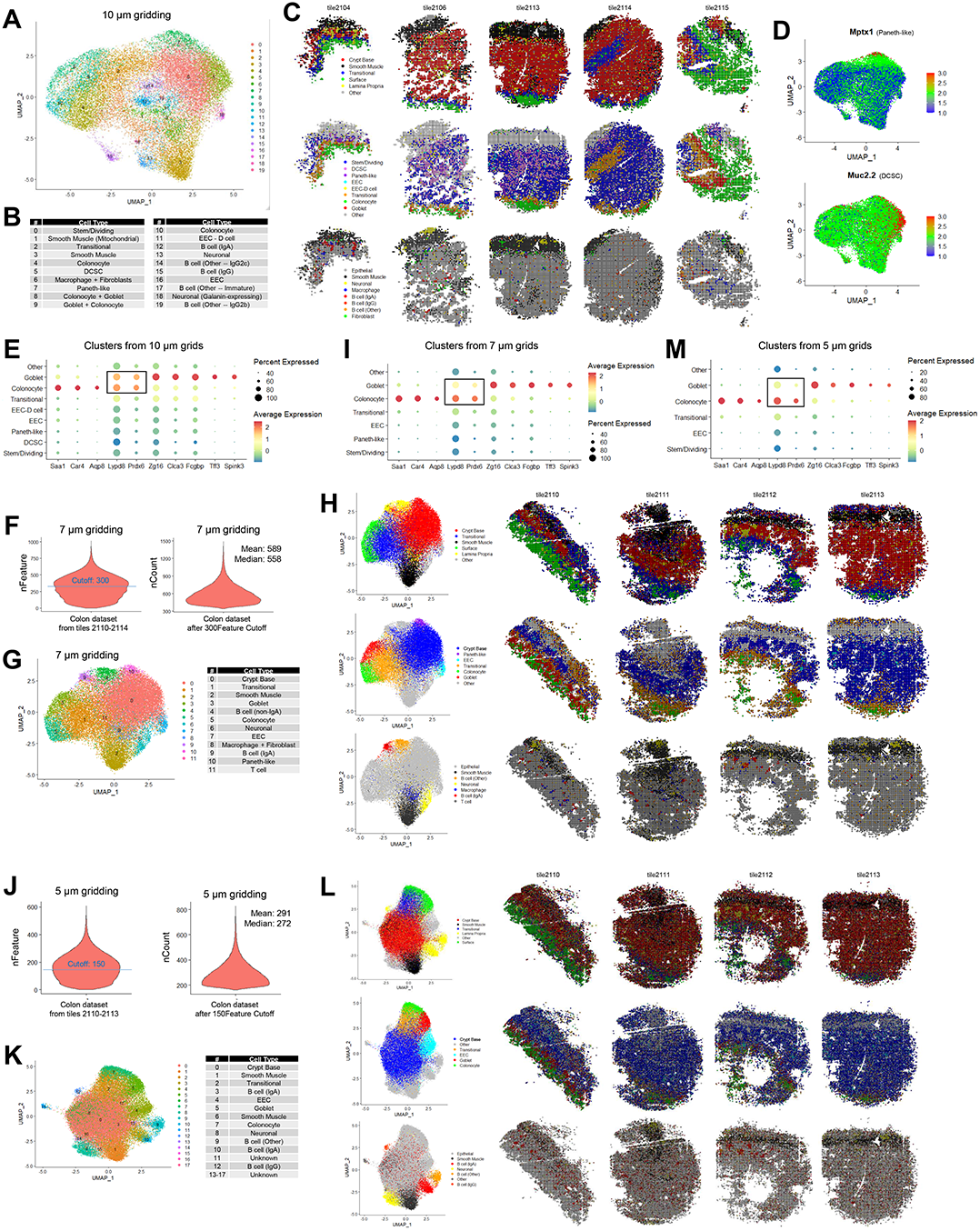
Seq-Scope Analysis of Colonic Spatial Transcriptome. (A-E) From the colon dataset binned with 10 μm square grids, a UMAP plot visualizing all clusters (A and B), spatial plots visualizing major histological layers (C, top; see Fig. 6B for cluster assignment), epithelial cell diversity (C, middle; see Fig. 6E for cluster assignment), and non-epithelial cell diversity (C, bottom; see Fig. 6I for cluster assignment), gene expression plots visualizing expression of *Mptx1* (D, upper; Paneth-like marker) and *Mup2* (D, lower; DCSC marker) in the UMAP manifold, and a dot plot showing colonocyte and goblet cell marker expression across different epithelial cell clusters (E) are presented. Box in (E) indicates that some colonocyte-specific markers, such as *Lypd8* and *Prdx6*, were found in Goblet cell grids, suggesting that colonocyte and goblet cell transcriptome is mixed in the Goblet grid pixel at this binning scheme. (F-I) Number of gene features (F, left; nFeatures) and UMI counts (F, right; nCounts after nFeatures cutoff at 300) were calculated across the indicated tiles of liver Seq-Scope dataset, binned using 7 μm square grids. From this dataset, a UMAP plot visualizing all clusters (G), UMAP plots (H, left) and spatial plots (H, right) visualizing major histological layers (H, top), epithelial cell diversity (H, middle) and non-epithelial cell diversity (H, bottom), and a dot plot showing colonocyte and goblet cell marker expression across different epithelial cell clusters (I) are presented. Box in (I) indicates that 7 μm square-gridded dataset performs better at separating colonocyte and goblet cell transcriptome compared to the 10 μm square-gridded dataset (E). (J-M) Number of gene features (J, left; nFeatures) and UMI counts (J, right; nCounts after nFeatures cutoff at 150) were calculated across the indicated tiles of liver Seq-Scope dataset, binned using 5 μm square grids. From this dataset, a UMAP plot visualizing all clusters (K), UMAP plots (L, left) and spatial plots (L, right) visualizing major histological layers (L, top), epithelial cell diversity (L, middle) and non-epithelial cell diversity (L, bottom), and a dot plot showing colonocyte and goblet cell marker expression across different epithelial cell clusters (M) are presented. Box in (M) indicates that 5 μm square-gridded dataset performs better at separating colonocyte and goblet cell transcriptome compared to the 10 μm (E) or 7 μm (I) square-gridded datasets.

**Fig S7.**
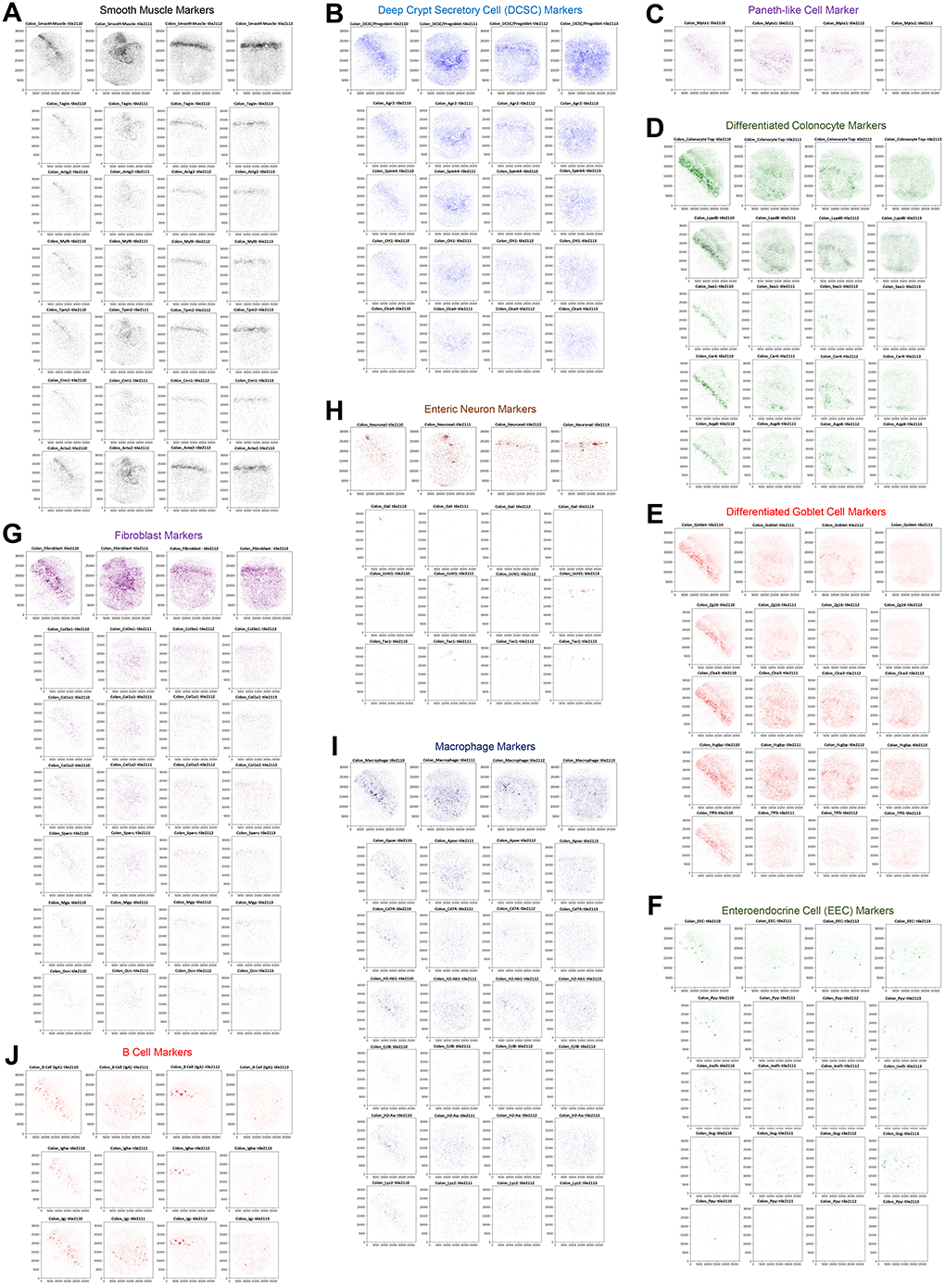
Seq-Scope Visualized Spatial Expression Patterns of Different Colonic Cell Type Markers. (A-J) Marker genes for indicated cell types were plotted onto the histological coordinate plane with indicated colors. Top row of each panel represents combined plotting of all listed markers. Bottom rows represent gene expression plotting of individual cell type marker genes. For all spatial plots, width and height of the imaging areas are 800 μm and 1 mm, respectively.

## Supplemental Table Legends

Table S1. List of genes that show cluster-specific expression patterns in the Seq-Scope liver dataset binned with square grids. p_val, unadjusted P value; avg_logFC, log fold-change of the average expression between the cluster and the background; pct.1, the percentage of cells where the gene is detected in the cluster; pct.2, the percentage of cells where the gene is detected in the background; p_val_adj, bonferroni-corrected p-values; cluster, the original cluster identity assigned by FindCluster function; Cell_Type, cell type identity manually assigned to the cluster.

(Tab 1: Normal 10 um) List of cluster-specific markers in the Seq-Scope normal liver dataset binned with 10 μm square grids. For UMAP visualization, please see Fig. S4A.

(Tab 2: Normal 7 um) List of cluster-specific markers in the Seq-Scope normal liver dataset binned with 7 μm square grids. For UMAP visualization, please see Fig. S4I.

(Tab 3: Normal 5 um) List of cluster-specific markers in the Seq-Scope normal liver dataset binned with 5 μm square grids. For UMAP visualization, please see Fig. S4O.

(Tab 4: TD 10 um) List of cluster-specific markers in the Seq-Scope *Tsc1*^*Δhep*^/*Depdc5*^*Δhep*^ (TD) liver dataset binned with 10 μm square grids. For UMAP visualization, please see Fig. S5A

Table S2. List of genes that show cluster-specific expression patterns in the Seq-Scope colon dataset binned with square grids. p_val, unadjusted P value; avg_logFC, log fold-change of the average expression between the cluster and the background; pct.1, the percentage of cells where the gene is detected in the cluster; pct.2, the percentage of cells where the gene is detected in the background; p_val_adj, bonferroni-corrected p-values; Layer, histological layer assigned to the cluster; Cell_Type, cell type identity manually assigned to the cluster.

(Tab 1: 10 um Original) List of cluster-specific markers in the Seq-Scope colon dataset binned with 10 μm square grids. For UMAP visualization, please see Fig. 6A and S6A.

(Tab 2: 10 um Epithelial) List of cluster-specific markers in the 10 μm square-gridded colon dataset with a refined epithelial cell type assignment (Fig. 6E).

(Tab 3: 10 um Non-epithelial) List of cluster-specific markers in the 10 μm square-gridded colon dataset with a refined non-epithelial cell type assignment (Fig. 6I).

(Tab 4: 7 um Original) List of cluster-specific markers in the Seq-Scope colon dataset binned with 7 μm square grids. For UMAP visualization, please see Fig. S6G.

(Tab 5: 5 um Original) List of cluster-specific markers in the Seq-Scope colon dataset binned with 5 μm square grids. For UMAP visualization, please see Fig. S6K.

Table S3. List of cell type markers that were used to generate spatial plots. Marker gene list used in each figure is summarized in each tab of the excel file.

## Notes

### Competing Interest Statement

Jun Hee Lee is an inventor on pending patent applications related to the development of Seq-Scope.

